# A real-time forecasting framework for emerging infectious diseases affecting animal populations

**DOI:** 10.1101/2024.12.12.628251

**Authors:** Meryl Theng, Chris Baker, Simin Lee, Andrew Breed, Sharon Roche, Emily Sellens, Catherine Fraser, Kelly Wood, Chris P. Jewell, Mark A. Stevenson, Simon M. Firestone

**Affiliations:** Melbourne Veterinary School, The University of Melbourne, Grattan Street, Parkville, 3010, Australia; Centre of Excellence for Biosecurity Risk Analysis, The University of Melbourne, Grattan Street, Parkville, 3010, Australia; School of Mathematics and Statistics, The University of Melbourne, Grattan Street, Parkville, 3010, Australia; Melbourne Centre for Data Science, The University of Melbourne, Grattan Street, Parkville, 3010, Australia; Epidemiology, Surveillance and Laboratory Section, Australian Government Department of Agriculture, Fisheries and Forestry, 70 Northbourne Avenue, Canberra, 2601, Australia; School of Veterinary Science, University of Queensland, Building 8114, Brisbane, Gatton 4343, Australia; Biosecurity & Food Safety, New South Wales Department of Primary Industries, 105 Prince Steet, Orange, NSW 2800, Australia; Department of Mathematics and Statistics, Lancaster University, Bailrigg, Lancaster, LA1 4YW, United Kingdom

**Keywords:** livestock disease, decision making, approximate Bayesian computation, spatiotemporal

## Abstract

Infectious disease forecasting has become increasingly important in public health, as demonstrated during the COVID-19 pandemic. However, forecasting tools for emergency animal diseases, particularly those offering real-time decision support when parameters governing disease dynamics are unknown, remain limited. We introduce a generalised modelling framework for near-real-time forecasting of the temporal and spatial spread of infectious livestock diseases using data from the early stages of an outbreak. We applied the framework to the 2007 equine influenza outbreak in Australia, generating prediction targets at three timepoints across four regional clusters. Our targets included future daily case counts, outbreak size, peak timing and duration, and spatial distributions of future spread. We evaluated how well the forecasts predicted daily cases and the spatial distribution of case counts, using skill scores as a benchmark for future model improvements. Forecast accuracy, certainty, and skill improved significantly after the outbreak’s peak, while early predictions were more variable, suggesting that pre-peak forecasts should be interpreted with caution. Spatial forecasts maintained positive skill throughout the outbreak, supporting their use in guiding response priorities. This framework provides a tool for real-time decision-making during livestock disease outbreaks and establishes a foundation for future refinements and applications to other animal diseases.

## 1 Introduction

Rapidly spreading infectious animal disease outbreaks pose serious risks to food security, international trade, livelihoods, animal welfare, and public health. Major examples include the 2001 foot-and-mouth disease (FMD) epidemic in the UK, the 2005 global high pathogenicity avian influenza H5N1 (HPAI) outbreak, 2007–2016 Q fever outbreak in the Netherlands, and the 2019–2021 lumpy skin disease (LSD) outbreaks in Asia. Responding to these emergencies is challenging for animal health authorities and livestock industries, as they must make critical, time-sensitive decisions based on an uncertain future. To help address this, mathematical modelling has become an important tool for guiding outbreak response, as shown during the COVID-19 pandemic, offering insights into what can be applied in animal health crises.

A range of mathematical models have been developed to support outbreak preparedness and response, varying greatly in complexity. On the simpler side of the spectrum, there are phenomenological approaches, such as the logistic growth model and the generalised Richards model, that make minimal assumptions about disease dynamics (e.g., Pell et al. 2018, Shanafelt et al. 2018). On the other hand, there are highly sophisticated simulation models that aim to recreate the behaviour of an outbreak and play out various response scenarios (e.g., Australian Animal Disease Spread Model or AADIS). The former is used for rapid estimation of key epidemiological parameters (e.g., infection rate and final outbreak size), while the latter tends to be used in longer-term strategic planning during “peacetime”, such as evaluating control strategies. Increasingly available computational power has expanded the types of models that can be subject to rigorous statistical analysis, leading to greater integration of the two modelling approaches and developments into data-driven approaches to mechanistic modelling (Lessler et al. 2016). The ability to fit time-series data to generative mechanistic models in a timely manner has enabled significant advancements in real-time forecasting, which has provided critical decision-support during recent public health crises (e.g., Viboud et al. 2018, Lutz et al. 2019, Moss et al. 2023).

While data-driven models for real-time forecasting are well-established and rapidly advancing in public health, similar tools for animal health are relatively limited and untested. A common modelling approach in animal health involves simulating an entire outbreak with defined parameters either fixed or sampled from distributions based on the published literature and expert opinion (e.g., AADIS, InterSpread). In contrast, more data-driven mechanistic approaches tend to be formulated with fewer parameters, mostly unknown, that are inferred from the data through a statistical fitting process. Jewell et al. (2009) made initial developments into this area with a full Bayesian Markov chain Monte Carlo (MCMC) approach to infer the underlying process and probability of undetected infections at premisess during the 2007 UK outbreak of FMD. Subsequent studies have adapted this method and include applications for: retrospective estimation of epidemiological parameters for the 2007 equine influenza (EI) outbreak in Australia (Firestone 2012); elucidation of spatiotemporal patterns of the HPAI H5N1 outbreak in the Mekong River Delta (Minh et al. 2011); and comparison between real-time and retrospective evaluation of control policies for the 2007 and 2010 FMD outbreaks in the UK and Miyazaki, Japan (Probert 2018).

Despite the potential of Bayesian MCMC approaches for generating risk predictions and forecasts, several limitations persist (Jewell et al. 2009). First, MCMC is a serial algorithm that cannot be trivially parallelised to enhance performance, making runtime a major issue. Second, we currently lack software to efficiently implement epidemic models, requiring new training algorithms for each structural change to the model. Third, since most transition events (i.e., from susceptible to exposed states) are unobserved, we must treat transition timings as a large, highly correlated, latent data space. This demands efficient sampling methods to estimate the posterior distribution accurately, but existing tools lack the speed and flexibility needed for real-time applications.

In addition to challenges with MCMC fitting algorithms, the performance of forecasts remains largely unexplored and poorly understood – particularly how forecast accuracy could vary across time, regions, and disease types, as a result of unmodelled mechanisms or data availability. These gaps highlight the need for more adaptable modelling frameworks that can quickly modify or compare different models during an outbreak without the need to recode likelihood functions (Drovandi et al. 2016). Addressing these limitations requires developing flexible and efficient approaches, alongside rigorous evaluation of their forecasting capabilities.

Here, we introduce a generalised modelling framework that produces near-real-time forecasts of temporal and spatial disease spread using data available in the first few weeks of a livestock disease outbreak and demonstrate how its predictions can be used to support decision-making during that period. We assume that existing premises- and herd-level data (location details, premises area and number of animals) are available prior to an outbreak, as we use this information to model how infections spread within and between these properties. As an outbreak progresses, we receive updated information on confirmed infectious premises (i.e., outbreak surveillance data), and this time series data is used to estimate parameters in our model using a likelihood-free Bayesian approach. Forward simulations (i.e., from the date of observation) generated from these parameter estimates represent the forecasts from the model. We apply our modelling framework on time slices of the dataset for the largest clusters of the 2007 EI outbreak in Australia, with the view of extending it to higher-priority, more complex emergency livestock diseases (e.g., FMD, LSD). We evaluate the model’s performance on this dataset by measuring forecast skill relative to a naive benchmark forecast, and in doing so, establish a baseline from which future extensions and improvements can be developed and evaluated against.

## 2 Methods

First, we discuss a generic modelling framework applicable to a broad range of animal disease epidemics (Figure 1).

**Figure 1:**
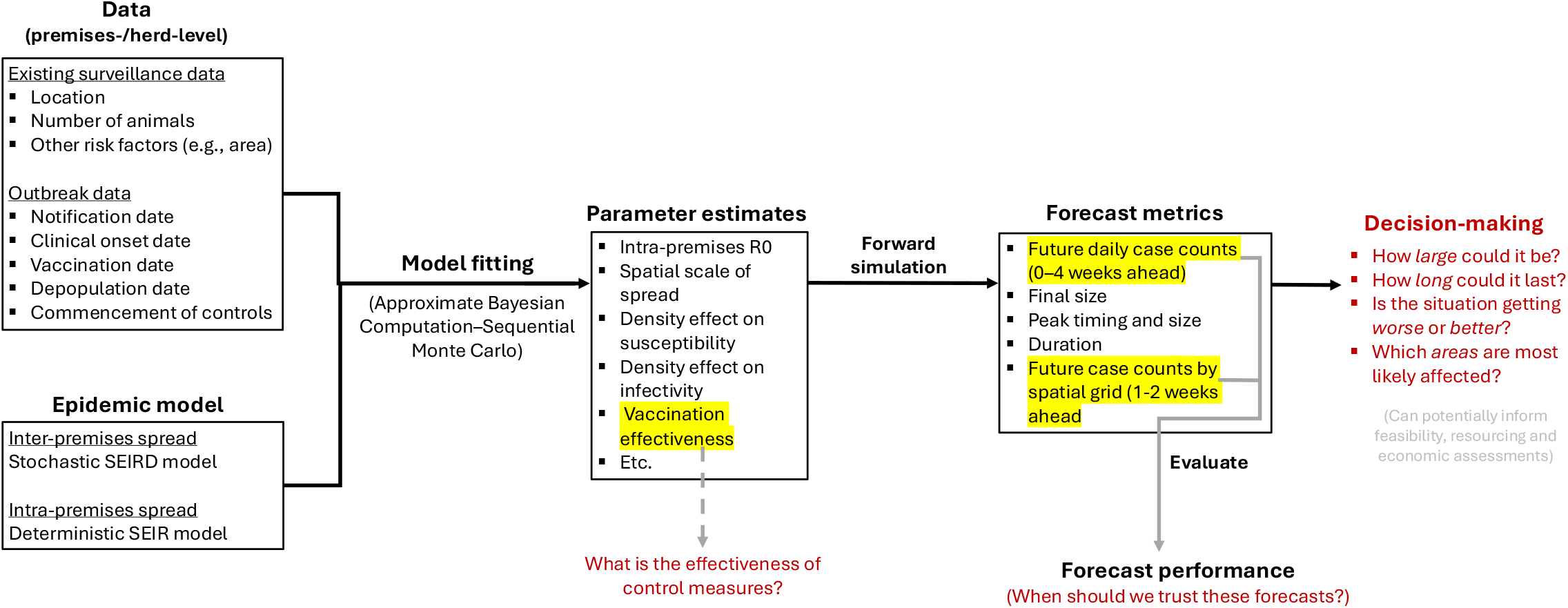
Conceptual representation of our modelling framework for real-time forecasting of a livestock disease outbreak. Model outputs can directly address questions (in red) to support decision-making, potentially informing feasibility, resource allocation, and economic assessments.

We begin by describing the epidemic model in Section 2.1, which comprises: (i) a stochastic SEIR (Susceptible, Exposed, Infectious, Recovered) model of inter-premises disease transmission influenced by premises area, number of animals and distance to other premises; and (ii) a deterministic SEIR model of intra-premises transmission between animals. We then outline an efficient approximate Bayesian computation (ABC) approach, in which we accept the simulations (i.e., particles) that are sufficiently matched to the observed data available at a given point in time (in Section 2.2). In Section 2.3, we characterise forward projections of the accepted particles (i.e., data simulated beyond the timepoint) with five forecast targets – (i) future daily case counts, (ii) final size of outbreak, (iii) peak timing and size, (iv) duration of outbreak, (v) spatial pattern of infection risk, (vi) future case counts by spatial grid – that focus on addressing the questions: “How large could the outbreak get?”; “Is the situation getting worse or better?”; and “Which areas/premises are most likely affected?”. We then detail an evaluation approach that measures how accurately a forecast predicted future daily case counts and future case counts by spatial grid relative to a naive bench-mark forecast, which addresses the question “When should we trust these forecasts?” (in Section 3.3).

To illustrate our methodology, we generated predictions by applying our framework to premises-level data from the 2007 equine influenza outbreak in Australia (in Section 2.4). We observed the epidemic at three time points (3, 5 and 7 weeks after the outbreak was first detected; Table 1), using the data available to produce forecasts for four of the largest outbreak clusters (Sydney Basin, Central-Coast-Maitland-Newcastle, Tamworth and Hunter Valley; see Section 2.4.1 for background). We show how forecast targets, parameter estimates and forecast performance refine as the epidemic progresses and how they vary between clusters.

**Table 1:** Descriptive statistics for highly affected regions (or clusters) in the Australian 2007 equine influenza outbreak, used as study sites in the development and validation of a real-time model of equine influenza in Australia.

|  | Sydney | CCMN | Tamworth | Hunter Valley | All outbreak affected areas |
| --- | --- | --- | --- | --- | --- |
| Number of horse premises | 7387 | 6549 | 2317 | 1522 | 103,221 |
| Number of infected premises | 2875 | 1293 | 582 | 404 | 8858 |
| Date of first detection | 25 August 2007 | 26 August 2007 | 26 August 2007 | 28 August 2007 | 24 August 2007 |

### 2.1 Epidemic model

The basis for forecasting is a stochastic model of the transmission mechanism between individual livestock premises (adapted from Jewell et al. 2009, Firestone 2012). Hence, the unit of interest is at the premises level rather the individual animal. We make two key assumptions in the model.

Firstly, we assume that out of a total population size of *N* premises, each premises is in one of four states at any given time point *t*:

- **S**usceptible premises do not have the disease and are able to be infected by it;
- **E**xposed premises have been infected with the disease but are not yet infectious themselves;
- **I**nfectious premises have been infected and are able to infect susceptible premises;
- **R**ecovered (or removed) premises that have either had all their animals recovered from the disease or culled, and therefore play no further part in the epidemic.

The only transitions between states allowed are: from susceptible to exposed, from exposed to infectious, and from infectious to recovered (or removed). Each infectious premises *i* was associated with event (i.e., transition) times *t*_*e*,*i*_, *t*_*i*,*i*_ and *t*_*r*,*i*_ representing the time of exposure, onset of infectiousness and recovery, respectively. Since these times are not (or tend not to) be directly observable, we have to impute them from observable data, such as time of onset of clinical signs, time of notification (to the appropriate animal health authority) and time of removal or sanitary measures (intended to render the premises no longer infectious). See Section 2.1.2 for how we can impute these event times.

Secondly, we allow for time-varying infectivity of premises, corresponding to intra-premises infection of more and more animals over time (Jewell et al. 2009). We model intra-premises infectivity using a deterministic SEIR compartmental model based on the number of animals recorded as present on each premises at the start of the outbreak (described in Section 2.1.2). The intra-premises transmission model is used to determine a premises’ infectious periods (i.e., recovery time) and the prevalence of infectious animals at time *t*, which contributes to premises-to-premises infection pressure (see Section 2.1.2).

#### 2.1.1 Transmission (inter-premises)

The inter-premises model is driven by a spatiotemporal inhomogeneous Poisson process, estimated with daily time-steps, using a tau-leap algorithm (Gillespie 1977, Jewell et al. 2009). Essentially, this is a spatially and temporally explicit modelling process that allows for different rates of new exposures to occur over time and space driven by the infection pressure in the system.

We start with the overall disease transmission rate, τ_*t*_, the sum of the individual infection pressures β_*ij*_ acting on all susceptible premises *j* from all infectious premises *i* at time *t*:

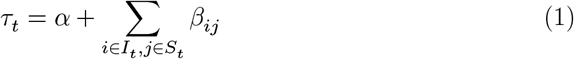

where α is the background transmission rate representing transmission not explicitly modelled (e.g., new seeding events from outside the population).

Pair-wise infectious pressure between individual infectious and susceptible premises β_*ij*_ was formulated as the product of three functions representing (i) the infectivity of premises *i*, (ii) the susceptibility of premises *j*, and (iii) the decay of infectious pressure with increasing Euclidean distance between *i* and *j*, giving:

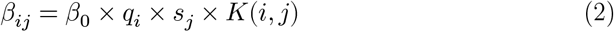

where β_0_ is the baseline transmission rate, *q*_*i*_ is the infectivity of premises *i* at time *t, s*_*j*_ the susceptibility of premises *j* at time *t*, and *K*(*i, j*) is a spatial kernel describing the transmission rate decay with increasing distance between *i* and *j*. These three functions can be formulated specific to each disease scenario and may be easily modified.

Here, we use a single-species formulation and defined the three functions as follows: First, the infectivity of premises *i* at time *t* is defined as,

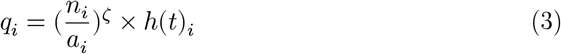

where *n*_*i*_ and *a*_*i*_ are the number of animals on, and area of infectious premises *i*, respectively, and ζ is an unknown parameter that allows for the non-linear effect of density of animals on infectivity. *h*(*t*)_*i*_ is the proportion of infective individuals within premises *i* at time *t* (i.e., prevalence), obtained directly from the solution of an SEIR system of differential equations which we describe in Section 2.1.2.

Second, the susceptibility of premises *j* at time *t* is defined as,

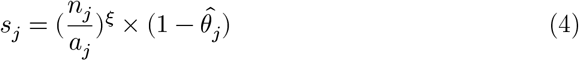

where *n*_*j*_ and *a*_*j*_ are the number of horses on, and area of infectious premises *j*, respectively, and *ξ* is an unknown parameter that allows for the non-linear effect of density of horses on susceptibility. The additional component, 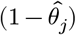, describes the effect of vaccination where immunity 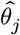 increases linearly from the date of vaccination *t*_*v*,*j*_ to full effectiveness *θ* over 14 days, giving:

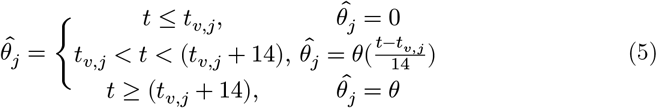

Finally, the spatial kernel *K*(*i, j*) was assumed to take the Cauchy form,

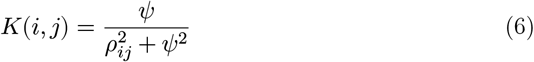

Where 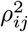 is the squared Euclidean distance between the centroids of premises *i* and *j*, and *ψ* is an unknown scaling parameter. The Cauchy form was selected for the spatial transmission kernel because it is computationally efficient, and it assumes a heavier tail (compared to exponential or geometric kernels) allowing for the possibility of long-range transmission. The spatial kernel can be selected from a range of options including power law and exponential decay functions, with the decision based either pragmatically on computational aspects or after testing for the best fitting form based on the data (Rorres et al. 2010, Boender and Hagenaars 2023).

#### 2.1.2 Infectious periods and prevalence (intra-premises)

The following differential equations are used to model the epidemic within each premises (herd, or other unit of interest), initialised for each newly infectious premises *i* at the time that it is simulated to be exposed *t*_*e*,*i*_, randomly sampling the number of initially exposed animals on the premises *e*,

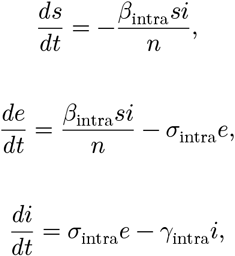

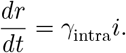

β_intra_, *σ*_intra_, and *γ*_intra_ represent the three transmission rates, and the counts of animals in each state are represented by *s, e, i* and *r* (totalling to *n*).

The prevalence of infectious individuals within a premises at time *t*, 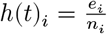, contributes to the calculation of infection pressures and the thinning of the Poisson process in the inter-premises model (Firestone 2012). Each newly infectious premises, *i*, is assumed to recover at time *t*_*r*_ when there is < 0.5 infections animals on the premises. Assumptions about the timing of onset of infectiousness (*t*_*i*,*i*_) and onset of clinical signs (*t*_*c*,*i*_) are scenario-specific and typically involve sampling from Gamma or Weibull distributions based on published details of the latent period (*t*_*i*,*i*_ −*t*_*e*,*i*_), incubation period (*t*_*c*,*i*_ − *t*_*e*,*i*_) and infectious periods (*t*_*r*,*i*_ − *t*_*i*,*i*_). Delay from onset of clinical signs to timing of notification (*t*_*n*,*i*_ − *t*_*c*,*i*_), and delay from notification to timing of depopulation (*t*_*d*,*i*_ − *t*_*n*,*i*_, where applicable) may be fixed at a single value, entered as a distribution to be stochastically sampled or or directly inputted for premises in specific regions at specific times. See Section 2.4.2 for details of the assumptions used for our case study.

### 2.2 Model fitting (ABC-SMC)

The reliability of forecasts generated by a model depends on how reasonably it was parameterised. Bayesian approaches are typically used for model-based statistical inference of unknown parameters in stochastic epidemic models. More recently, approximate Bayesian computation (ABC) methods have gained traction in infectious disease modelling (e.g., Minter and Retkute 2019). Unlike MCMC methods, ABC approximates parameter posterior distributions without likelihood functions, which can be intractable or computationally costly to evaluate (Beaumont 2010). The flexibility of ABC approaches provides an attractive way to fit models to often incomplete and/or complex data in an accurate and timely manner.

We fit model outputs to observed data (daily case counts and total grid-cell case counts), using ABC with a particle filter algorithm called sequential Monte Carlo (ABC-SMC) (Beaumont 2010). Like all ABC methods, this involves sampling a candidate set of parameter values, *θ*^*^ from the prior distributions *P* (*θ*), and using this to simulate data from a model (*D*^*^), in this case representing a time series of daily case counts and total grid-cell case counts. These are compared to the observed data *D* by using a distance function *d*(*D, D*^*^), and the candidate values satisfying *d*(*D, D*^*^) < ϵ for some threshold ϵ are ‘accepted’. This process, known as ABC rejection sampling, repeats until *N*_px_ ‘particles’ (i.e., parameter sets) are accepted, providing a set of particles that form posterior distributions of model parameters. However, naive rejection sampling is very inefficient because it explores the entire parameter space. ABC-SMC improves efficiency by propagating the set of particles through a sequence of intermediate distributions by gradually decreasing tolerance ϵ_1_ > ϵ_2_ > …, > ϵ_*G*_. Each intermediate distribution (called a generation) is obtained as a weighted sample from the previous distribution that has been perturbed through a kernel *K*(*θ*|*θ*^*^).

We adapt an ABC-SMC algorithm described by Minter and Retkute (2019) to our two distance functions, one temporal and one spatial. To account for reporting delays, we only consider data starting from the day of first detection in a cluster (*t*_detect_ = earliest notification) and ending at the timepoint minus some number of days *T* = *t*_today_ −*t*_lag_.

The temporal distance function is the sum of absolute differences of daily case counts (number of exposed premises per day *E*_*t*_) between the observed and simulated data,

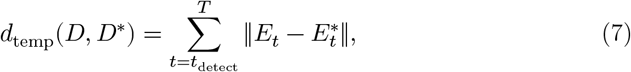

for *t* = *t*_detect_, …, *T* .

The spatial distance function is the Pearson’s correlation coefficient between the grid-specific total case counts (*E* = (*E*_1_, *E*_2_, …, *E*_𝒳_)) of observed and simulated data for *t* = *t*_detect_, …, *T*,

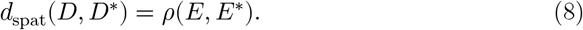

Here, the spatial extent of the outbreak cluster is represented by 𝒳 = *X* × *X* grid cells, and grid cells that do not contain premises are excluded.

We accept *θ*^*^ if the temporal distance is less than or equals to the temporal tolerance (*d*_temp_(*D, D*^*^) ≤ ϵ_temp,*g*_) *and* the spatial distance is more than or equals to the spatial tolerance for generation *g* (*d*_temp_(*D, D*^*^) ≥ ϵ_spat,*g*_).

We used a normal distribution as our perturbation kernel following Beaumont (2010), where 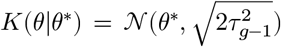, and the weight denominator for *g* > 1 is 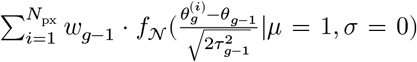, where *f*_𝒩_ is the probability density for the normal distribution and 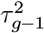 is the weighted empirical variance of the 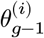 s. Details of the ABC-SMC algorithm used are provided in Appendix A.

### 2.3 Forecast targets and evaluation

#### 2.3.1 Targets

Forecast targets are the outcomes being predicted. Our model provides the following targets:

i. **future daily case counts:** daily case counts (exposed number of premises per day, ***E***_*t*_) for the 3 days prior to the timepoint, for the timepoint itself, and for the 4 weeks (28 days) after the timepoint
ii. **final size:** total case count over the course of the entire outbreak
iii. **peak timing and size:** day with the maximum case count and the case count for that day
iv. **Duration:** first day to the last day (with a non-zero case count) of the outbreak
v. **spatial pattern of infection risk** (1-2 weeks ahead and up to predicted end of outbreak): the risk that a given premises becomes infectious during the current outbreak, presented as a map; risk was estimated by the proportion of simulations (out of 1000) in which a presently susceptible premises was infectious between the date of first detection and (a) 1-week ahead from the current date, (b) 2-weeks from the current date, and (c) the end of the outbreak.
vi. **spatial distribution of future case counts** (1- and 2-weeks ahead): spatial grid-cell case counts (exposed number of premises per cell, ***E***_*x*_) between the 3 days prior to the timepoint and 1- or 2-weeks ahead from the timepoint.

#### 2.3.2 Evaluation

Forecast performance of the first and last target (i.e., future daily case counts, future grid-cell case counts) was quantified using Continuous Ranked Probability Scores (CRPS). CRPS is a widely-used metric for probabilistic forecasts (Gneiting and Katzfuss 2014). It penalises forecasts for placing probability mass at a distance from the ground truth, with values increasing as more probability mass is allocated to incorrect values, and as the distance between these incorrect values and the ground truth increases.

For the future daily case counts, we calculated CRPS values for each daily observation of case counts as per Moss et al. (2023). We then compared the performance of our model ℳ to that of a naive benchmark forecast 𝒩, which was simply the most recently observed value:

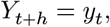

for *h* ∈ {−3 − 2, …, 27, 28}.

Forecast skill was calculated using the CRPS values for the model predictions (CRPS_ℳ_) and that of the naive benchmark forecast (CRPS_𝒩_):

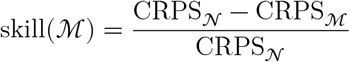

Negative skill scores indicate that the forecasts performed worse than the naive benchmark, zero indicates that the forecasts performed as well as the naive benchmark, and positive values indicate that the forecasts performed better than the naive benchmark. The maximum possible skill score is one, which indicates that a forecast was perfectly accurate with no uncertainty.

Similarly, for future grid-cell case counts, we calculated CRPS values for each observation of total case counts within a spatial grid cell for two periods: 1-week ahead (lead time of −3-7 days) and 2-weeks ahead (lead time of -3–14 days). Our naive benchmark forecast was the observed value for the most recent period of the same length:

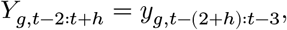

for *h* = 7 or 14.

### 2.4 Application: 2007 Australian equine influenza outbreak

#### 2.4.1 Background

Equine influenza is a highly contagious respiratory disease of horses. Australia experienced its first ever EI outbreak in August 2007, which spread rapidly throughout the largely unvaccinated horse population, infecting a reported total of 67,000 horses on 9,359 premises. Eradication was achieved in less than five months, through a combination of horse movement restrictions, on-premises biosecurity measures and targeted vaccination (Callinan 2008, Glanville and Christie 2011). Since then, numerous analyses of the 2007 outbreak have been conducted and datasets prepared for epidemiological analysis are available (Garner et al. 2011, Moloney 2011, Firestone 2012). The relatively simple and well-understood outbreak, along with the availability of suitable data and forecasting methods, presents an opportunity to simulate the process of providing real-time forecasts to decision-makers during an animal health crisis and to test the reliability of such forecasts.

In order to analyse this epidemic, we used a dataset that was prepared from premises-level covariate and contact tracing data provided by the New South Wales (NSW) and Queensland (QLD) State governments (data collation and cleaning details described in Cowled et al. 2009, Garner et al. 2011, Firestone 2012). These data contained the following covariates: address, geographical location (longitude and latitude of the centroid of the premises), number of horses present, premises area, vaccination status (and vaccination date for NSW data). Infectious premises data also included the estimated date of onset of first clinical signs of the first horse affected (‘onset date’).

The Sydney, Central Coast-Maitland-Newcastle, Tamworth and Hunter Valley regions were the four most severely affected clusters in the country, covering 89% of all infectious premises (Figure 2). These clusters were as described in (Firestone et al. 2011) with precise delineatation implemented in the Geographic Information System QGIS (QGIS Development Team 2024) following natural geographic boundaries and purposefully ensuring separation of neighboring clusters. Descriptive statistics for each cluster are listed in (Table 1).

**Figure 2:**
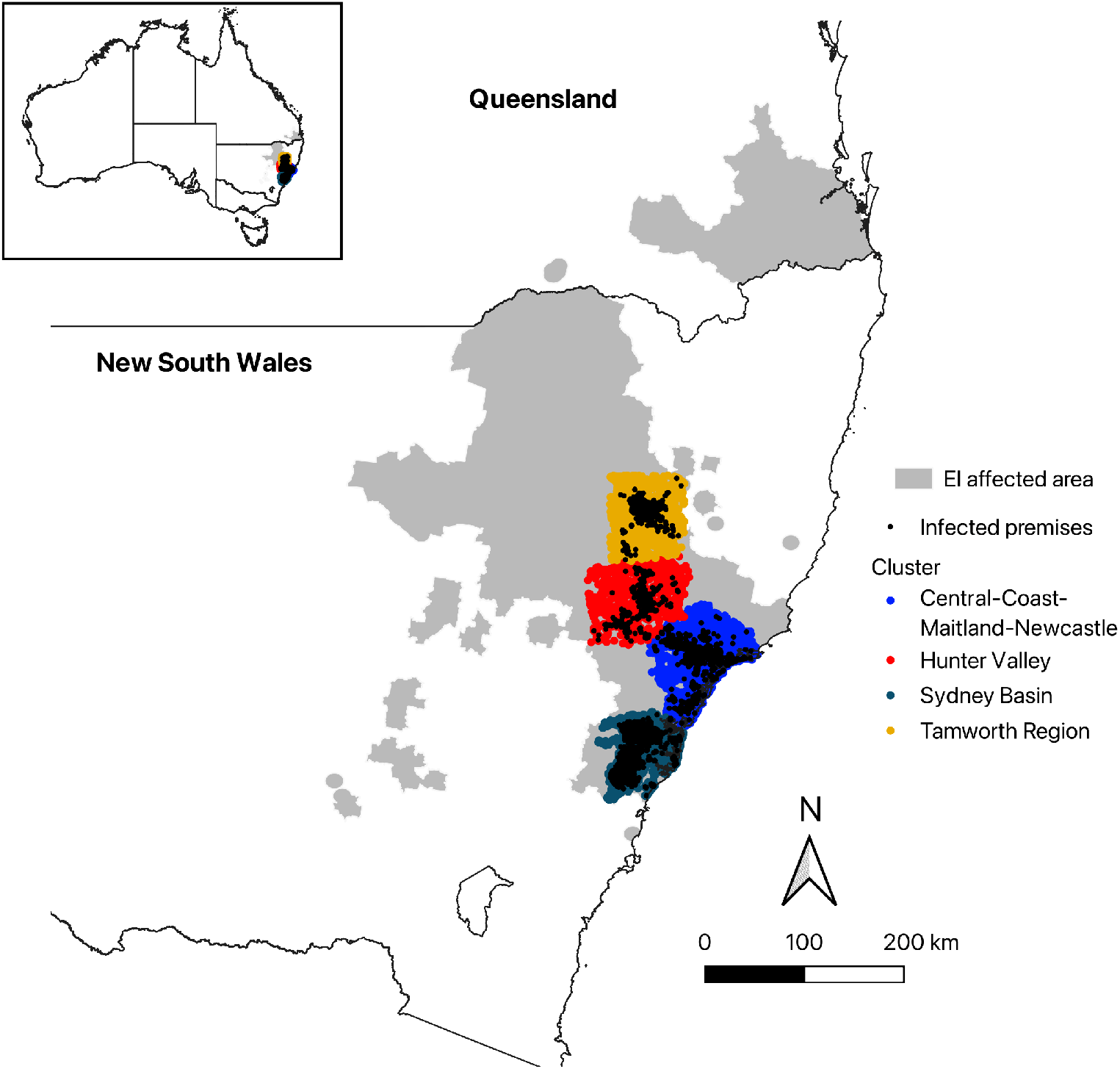
Four highly affected regions (or clusters) in the Australian 2007 equine influenza outbreak, used as study sites in the development and validation of a real-time model of equine influenza in Australia.

#### 2.4.2 Specific modelling details

##### 2.4.2.1 Transmission model

For each region and timepoint, our model was fitted to the EI dataset and the ABC-SMC algorithm was used to infer the posterior probability distribution for seven unknown parameters (α, β_0_, *ψ, ξ*, ζ, *θ*, β_intra_). Naive priors were placed on all unknown parameters.

The within-premises latent period 1/*σ*_intra_ was fixed at 1-day, and the recovery rate 1/*γ*_intra_ was fixed to 6-days based on experimental evidence (Mumford et al. 1990, Daly et al. 2011). Though, note that these can be treated as unknown parameters to be inferred by the model.

We derived exposure time *t*_*e*,*i*_ from the assumption that the incubation period for each premises (i.e., the time between exposure and clinical onset; *t*_*c*,*i*_ − *t*_*e*,*i*_) followed a Gamma distribution with a fixed shape parameter α = 4 and a rate parameter λ = 2, giving (*t*_*c*,*i*_ − *t*_*e*,*i*_) ∼ Gamma(4, 2). These parameters were selected to return a distribution that reflects the typical 1–3 day incubation period of equine influenza (Paillot et al. 2006), with a small probability of longer incubation periods (4–6 days) as observed in ponies experimentally infectious with low doses of virus (Mumford et al. 1990).

Finally, we fixed the reporting delay parameter *t*_lag_, which specifies the number of days before the current timepoint to fit the model up to, at 3 days.

##### 2.4.2.2 ABC-SMC

The number of desired particles was set as *N*_*px*_ = 1000 and the number of generations to *G* = 8 in all runs. For the largest cluster (Sydney Basin), the temporal tolerance value was set to allow an average distance of 50 cases per day for the first generation (ϵ_temp,1_ = 50 · (*T* − *t*_detect_ + 1)), and the value for each subsequent generation was progressively reduced, with the tolerance for the last generation set to six cases per day (ϵ_temp,8_ = 6 · (*T* − *t*_detect_ + 1)). These values were scaled based on the number of premises in the cluster relative to Sydney Basin. The spatial tolerance schedule was set to allow a minimum correlation coefficient of 0.3 in the first generation (ϵ_spat,1_ = 0.3) and progressively increasing to 0.725 in the last generation (ϵ_spat,1_ = 0.725).

Tolerance schedules were set based on pilot runs to balance the tightness of fit with efficiency of sampling model parameters. Analyses exceeding a run-time of two days (including computing cluster job queues) were terminated, and the particles from the latest completed generation (with *N*_*px*_ ≥ 400) formed our model output.

All parameter values and priors used for the transmission model and the ABC-SMC algorithm can be found in Table A1 in the Supplementary Material.

## 3 Results

We successfully fitted our model to each of the four clusters of the EI outbreak in Australia. For each cluster we fitted data to three timepoints of the outbreak (at 3, 5 and 7 weeks after first detection), to gain an understanding of forecast accuracy for different amounts of data. For each forecast, we produced the following forecast targets: (i) future daily case counts, (ii) final size of outbreak, (iii) peak timing and size, (iv) duration of outbreak, (v) spatial pattern of infection risk, and (vi) future grid-cell case counts. We assessed the quality of the forecasts at different timepoints by quantifying how well each forecast estimated future daily case counts up to 4-weeks ahead.

Overall, results showed that uncertainty recedes over the course of the epidemic. Generally, all targets provided good estimates with sufficient data, typically from the second timepoint (after 5-weeks), which tended to be the time when the peak in daily case counts had just been formed (Figure 3). At the first timepoint (3-weeks’ worth of data), estimates were much more variable.

**Figure 3:**
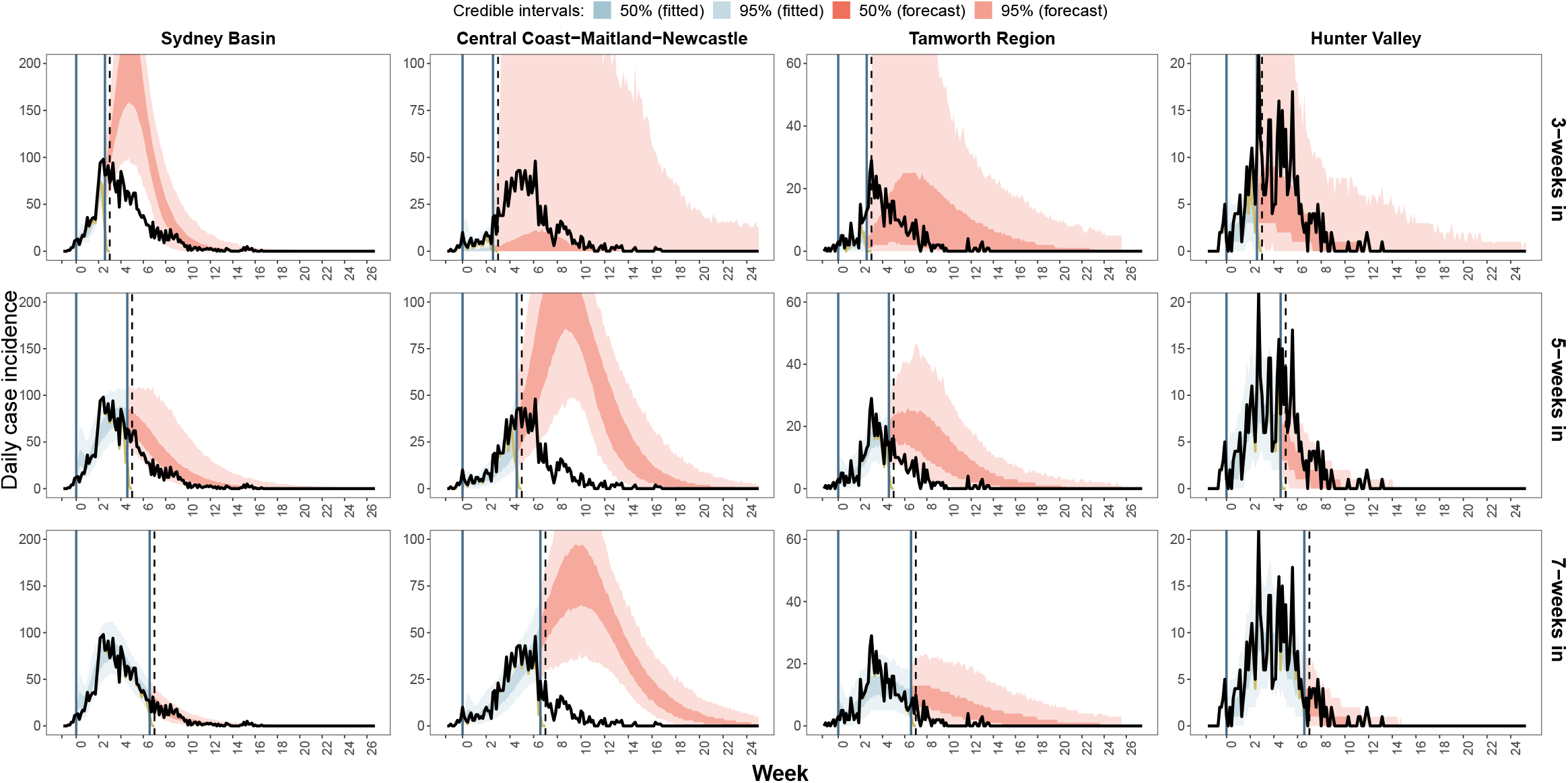
Forecast of daily EI case incidence for four outbreak clusters in Australia, shown separately for each time point (right-hand axis labels). The blue shaded regions represent model fit to the data available at the time of forecasting (dark yellow lines). The red shaded regions represent the forecast. Vertical black dashed lines indicate the time point. Vertical blue lines indicate the time window of data the model was fit to (accounting for delay with a 3-day lag). Black lines represent all reported daily case counts (i.e., all observations by end of outbreak). Shaded areas indicate the 50% and 95% percentile of 1000 simulated trajectories (of accepted particles).

In the following, we present the outputs generated by our modelling framework and demonstrate how they can address key questions to support decision-making. Posterior distributions of model parameters can be found in Appendix B.4.

### 3.1 Outbreak severity and trend

Forecasts generated at the first time point (3-weeks in or 21 days after first detection) were highly uncertain across all regional clusters other than Sydney, including the possibility that the outbreak *might* be negligible (i.e., 95% CrIs include possibility where <7% of premises are infectious) and the possibility that a large outbreak *might occur* (i.e., 95% CrIs include possibility of >69% of premises infectious) (Figure 3) (Table B1 in Appendix B.1). The forecast for Sydney, the largest cluster consisting 7387 premises, was the most precise, demonstrating confidence that the outbreak was accelerating (95% CrI trajectories with increasing daily case counts) up to a peak of 230 (95% CrI: 121–382) cases per day at end of Week 6 (day 42), and eventually infecting 56–99% (95% CrI) of premises within the region.

By the second time point (5-weeks in), forecasts across all clusters were more certain. Forecasts for the Sydney, Tamworth and Hunter clusters demonstrated confidence that the peak of the outbreak was near or had been reached. Forecasts for these three clusters were also in agreement with future case counts (95% CrIs captured the future case counts; albeit within the lower bounds for Sydney and Tamworth). At this time point, the observed peak of the outbreak just started to form for these clusters, but not in the CCMN cluster, where the forecast indicated that a larger outbreak *would occur* (50% CrI included trajectories with increasing case counts), peaking between 7–12 weeks and potentially infecting 60–94% of all premises (Table B1). The forecast for this region was a significant overestimation of the size of the outbreak.

As the peak of the outbreak became more established for the Sydney, Tamworth and Hunter clusters, subsequent forecasts (7-weeks in) yielded more confident and accurate predictions. In the CCMN cluster, where the peak of the outbreak was still not apparent at this time point, its forecast still demonstrated confidence that a growing outbreak *would occur*. Though forecasts beyond the observed peak for this cluster no longer predicted a much larger outbreak, the model struggled to fit the peak and predicted a relatively smaller but protracted outbreak (Appendix B.2).

### 3.2 Areas of immediate concern

To address the question “Which premises are likely to be affected?”, we consider the risk that a given premises becomes infectious during the current outbreak. This was estimated by the proportion of simulations (out of 1000) in which a presently susceptible premises was infectious between the date of first detection and (i) 1-week ahead from the observation date (i.e., *t*_today_), (ii) 2-weeks from the observation date, and (iii) the end of the outbreak. These are shown as spatial risk maps in Figure 4 and Appendix B.3, demonstrating how the risk estimate progresses with time within each cluster, and how they compare to the observed outbreak.

**Figure 4:**
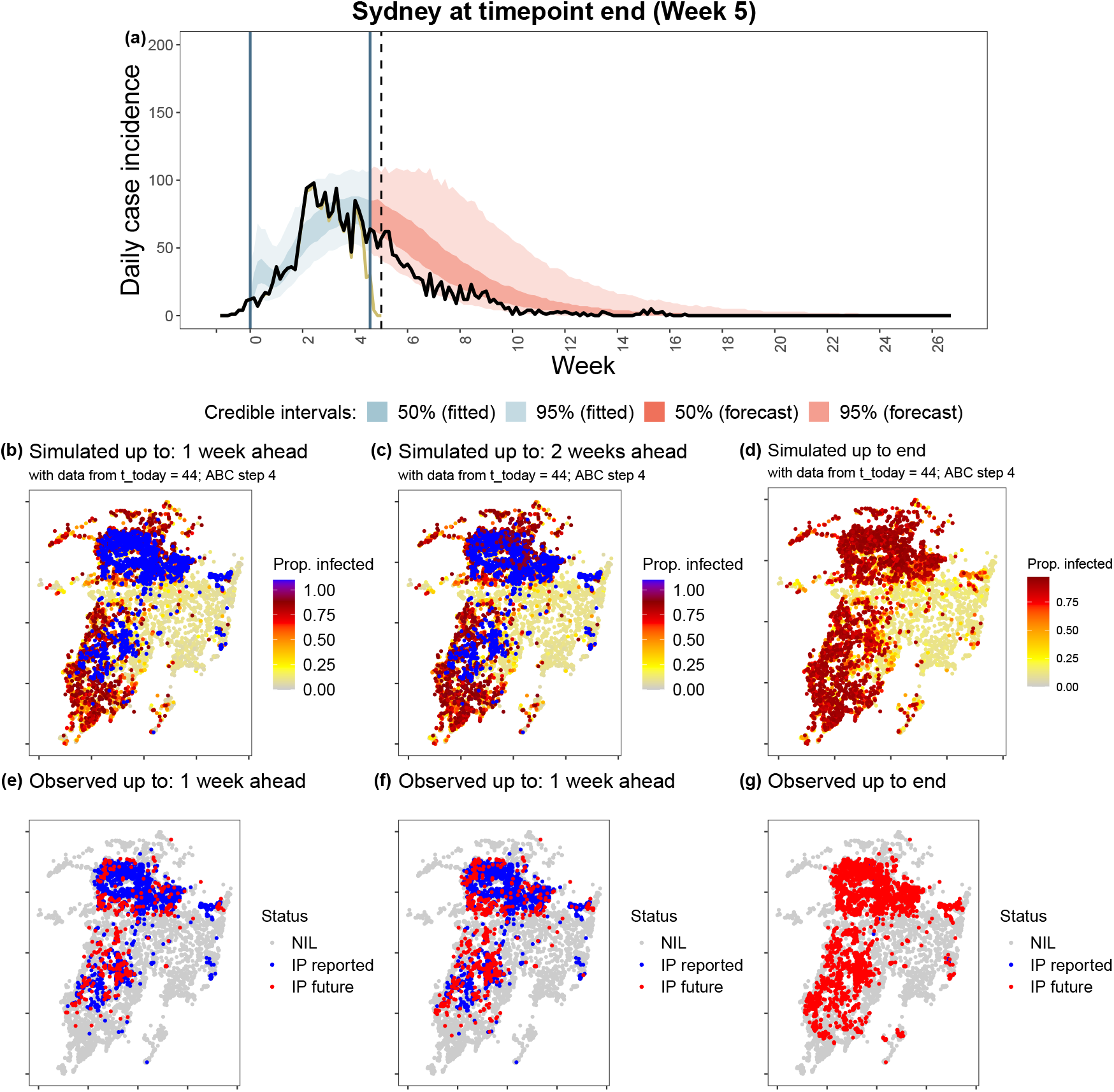
Comparison of simulated and observed outbreaks for the **Sydney Basin region at timepoint 2 (week 5).** (a) Simulated epidemic curve, with 50% and 95% credible interval (shaded regions), compared to all observations (black line) and to only observations available at the time of forecasting (dark yellow line). Spatial maps showing proportion of runs (out of 1000 simulation runs) in which a premises was infectious between the date of first detection and (b) 1-week ahead from the current date, (c) 2-weeks ahead from the current date, and (d) the end of the outbreak; compared to (e-g) the single observed realisation of the outbreak at the same respective dates (“NIL”: non-infectious premises; “IP reported”: infectious premises reported before the timepoint; “IP future”: infectious premises reported after the timepoint). Plots for other regions and timepoints can be found in Appendix B.3.

Spatial predictions at the first time point (3-weeks after first detection) tended to be too imprecise to provide a useful indication of risk areas across all clusters. From the second time point onward (5- and 7-weeks), spatial forecasts for each cluster provided clear indication of high risk areas in the next 1-2 weeks. Spatial risk predictions of 1- and 2-week ahead infection risk successfully captured general patterns of ‘future’ transmissions (in terms of relative risk) in all clusters. In Sydney, however, 1- and 2-week ahead, up-to-end forecasts predicted more widespread outbreaks (where ‘future’ cases tended to occur further away from already-reported cases; Figure 4). In CCMN, where outbreak severity was over-predicted, a much more widespread transmission was predicted up to the end of the outbreak.

### 3.3 Forecast performance

#### 3.3.1 4-weeks ahead daily case counts

Overall, forecast skill was higher for shorter lead times across all four outbreak clusters (Figure 5). The forecasts out-performed the naive benchmark (i.e., the most recently observed case count – 3-days before the day of prediction) for 0.5 and 1-week lead times in the Sydney and Tamworth clusters, respectively. Forecasts out-performed the naive benchmark for almost all lead times for the smallest cluster, Hunter Valley, which reported a maximum of 17 cases a day. The naive benchmark out-performed the forecasts for lead times beyond 0 weeks in CCMN, where the most recently observed case count was a good predictor of the future case counts.

**Figure 5:**
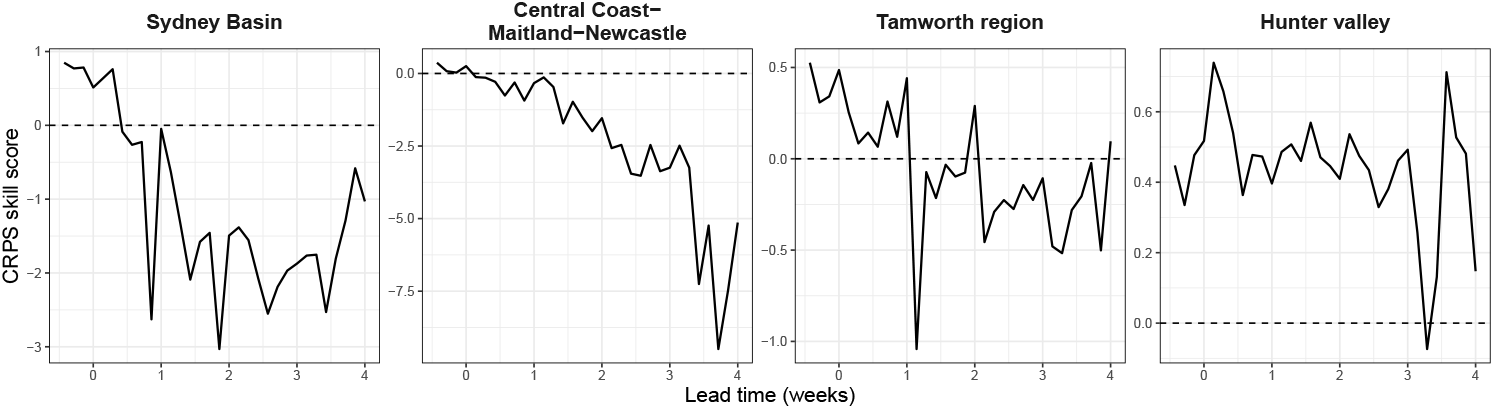
Forecast skill of EI daily case counts by lead time for four outbreak clusters in Australia, averaged across all three timepoints. Positive skill scores indicate that the forecasts outperformed the naive benchmark, with a maximum possible score of one (i.e., perfect accuracy and no uncertainty). Negative skill scores indicate underperformance relative to the naive benchmark, with a mininum score of −∞.

For longer lead times, the most recently observed case count – which under-reported cases ultimately recorded for the day (even with a 3-day lag) – better aligned with future observations showing a decrease in cases, and increased their skill relative to the model’s forecasts.

Forecast skill across all clusters was highest when daily case counts were 0 or ≥ 30, and was lower for days where 1–9 cases were reported (Figure B23).

##### 3.3.2 1-2 weeks ahead grid-cell case counts

For all clusters and timepoints, both the 1- and 2-weeks ahead spatial forecasts (of grid-cell case counts) outperformed the naive benchmark on average (Figure 6); except in one scenario (i.e., 2-weeks ahead forecast for Sydney Basin at the end of Week 3).

**Figure 6:**
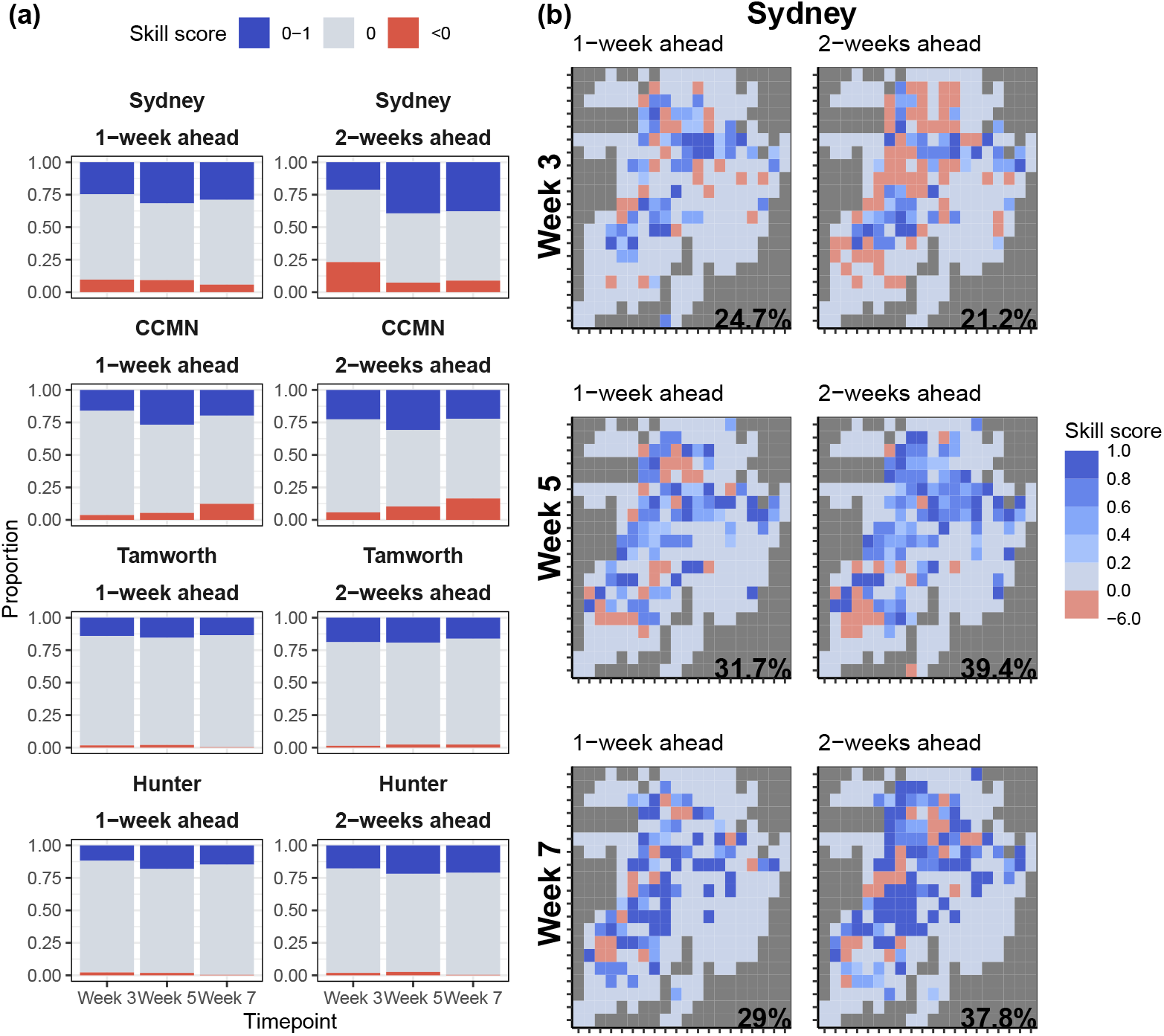
Spatial forecast skill for 1- and 2-week ahead EI case counts. (a) Summary of proportion of grids where the forecast outperformed (CRPS skill score of >0), performed the same (=0), or underperformed (<0) the naive benchmark. (b) Spatial distribution of forecast skill for 1- and 2-week ahead case counts for the Sydney Basin (see Figure B20 for other clusters).

Forecasts for the two largest clusters (Sydney Basin and CCMN) demonstrated the greatest proportion of area with positive skill over the naive forecast (i.e., out-performance), but also a greater proportion of area with negative skill over the naive forecast (i.e., under-performance) compared to that of the smallest clusters (Tamworth and Hunter Valley). Spatial forecasts for the Sydney Basin improved through the timepoints (i.e., increase in area of positive skill and decrease in area of negative skill). However, the difference in skill between timepoints in the other clusters was marginal (Figure B24).

## 4 Discussion

In this work, we introduced a real-time forecasting framework that can be generalised to any emergency infectious disease of livestock. Application to premises-level data from the 2007 EI outbreak in Australia demonstrated its potential for providing timely and effective spatiotemporal forecasts for different amounts of data and in different regions/clusters. Our initial evaluation of forecast performance shows that while forecast performance improved with time (i.e., due to more data accumulated), performance varied between clusters. We use findings from our case study to identify potential directions for further investigation, with a view to continuously improve forecast performance and extend application to other animal diseases in future work.

### 4.1 Principal findings

Our evaluation demonstrated that forecast performance generally improved throughout the outbreak. Forecast *uncertainty* reduced with time as case data increased in volume, which enabled more precise estimation in parameters describing disease spread (see Appendix B.2). Early in the outbreak (3-weeks), uncertainty tended to be high when reported caseloads up to the time of forecasting was low, as seen in the three smaller clusters where daily case incidence was below 10 at the first timepoint (3-weeks). Prediction uncertainty is a common problem early in the course of an epidemic (Castro et al. 2020, Wilke and Bergstrom 2020), a critical time during which decisions to intervene tend to be the most consequential. It is therefore crucial to incorporate all expert knowledge into prior distributions for parameters from the outset (Jewell et al. 2009), which can improve forecast precision in the face of scarce data. Forecast *accuracy*, on the other hand, improved with time as the impact of control measures became evident in the case data. In particular, the early movement ban enacted once the first EI case was detected was known to have substantially reduced the background spread rate (Firestone 2012). Hence, forecasts made before the peak in daily case counts had formed tended to overpredict outbreak severity (i.e., Sydney Basin at 3-weeks, CCMN at 5- and 7-weeks), likely representing what could have happened in the absence of movement restrictions. Explicit modelling of movement restrictions can be explored in future work, though this requires timely and specific policy information – i.e., the time and extent to which they are imposed or relaxed within each jurisdiction. We did, however, model vaccinations, which had a limited effect on the outbreak, likely due to their late implementation (see Appendix B.6 for details).

We also found that forecasts projecting closer into the future (i.e., shorter lead times) exhibited greatest skill, which suggests that greater trust be placed on nearer-term projections. This agrees with the general expectation that forecasts typically perform worse as they predict further ahead (Tong 1990), and is supported by both theoretical investigations (e.g., Castro et al. 2020) and empirical evidence from other outbreaks (Viboud et al. 2018, Scarpino and Petri 2019). Thus, it may be more useful to make decisions based on nearer-term predictions (e.g., 1-2 week ahead daily case counts) rather than on summary targets that attempt to characterise the whole outbreak (e.g., final size, duration, peak timing and size).

### 4.2 Study strengths

A key strength of our modelling framework is the ability to provide reliable spatial forecasts throughout the outbreak. Forecasts of 1-2 week ahead grid-cell case counts outperformed the naive benchmark despite several factors: the use of a simple/crude ABC-SMC fitting algorithm, uncertainty in the spatial kernel parameter estimates, and (in some scenarios) over-prediction in daily case counts. While forecasts of daily case counts did not perform as well as spatial forecasts overall (in terms of beating the naive benchmark), especially at the earliest timepoint, the latter is arguably much more useful in decision-making as it provides spatial information on high-risk areas that could be prioritised for outbreak control and surveillance.

Another strength of our methodology is the use of an ABC-SMC fitting approach for its generalisability to different outbreak contexts and computational efficiency. The two distance metrics used to fit the epidemic model to the data (i.e., distance between observed and simulated data in terms of daily case counts and grid-cell case counts) were simple, yet effective, and are generalisable to any model adaptation. Furthermore, our analysis generated forecasts in near-real-time, taking up 1-2 days to complete for the largest EI cluster (no. premises = 7387) on a computing cluster of 50 nodes (including job queue wait times). These features contrast with full MCMC approaches that have been used in similar modelling frameworks (e.g., Jewell et al. 2009, Firestone 2012), which can be much slower to program and perform with repeated calculation of the likelihood function (McKinley et al. 2009).

### 4.3 Limitations and challenges

In our initial attempt at configuring the framework for our EI case study, we prioritised model parsimony to suit a rapid model fitting process and to suit the data available. As such, there are several factors that we have not considered here. First, we assumed that background/baseline spread rates do not change over time. Time-dependent background or baseline spread rates are often used to account for the impact of interventions (e.g., Dehning et al. 2020, Moss et al. 2023). However, inference of time-varying rates as unknown parameters requires assumptions on future spread rates or between-premises movement data to inform these rates (e.g., Jewell et al. 2009), which we did not have for the EI outbreak. Secondly, there may some level of misspecification bias in our model formulation of factors driving disease spread as parameter sensitivity varied a great deal (see Appendix B.2). Alternative model formulations should be explored in future work and could be accompanied by sensitivity analyses as a guide. Our study serves as a benchmark from which alternative formulations can be evaluated against using the skill scoring approach we have developed. Third, we corrected for delays in case reporting by starting the forecast three days prior to the timepoint (i.e., *t*_lag_ = 3), which appeared to be sufficient in mitigating the effect of reporting delays on forecast performance in our case. However, this simple correction for delays will likely not apply in many situations, where often, delays between symptom onset and case reporting are context specific and are modelled based on available real-time data (e.g., Moss et al. 2023). Accounting for partial observations (e.g., right-censoring) can readily be done in our modelling framework and should be a priority in future work.

Apart from modelling decisions that may have limited model performance, we also noticed issues in our underlying data. We found location information for a number of horse premises to be inaccurate, which is a common problem across animal health surveillance data (Sheffield et al. 2018). This would have affected estimation of transmission distance and horse density (as premises area was estimated from locations, Firestone 2012), and consequently, parameter estimation and forecast reliability. Thus, regular updates and verification of livestock industry databases is critical, which can be assisted by emerging machine learning approaches (van Andel et al. 2017, Sheffield et al. 2018).

### 4.4 Implications and future work

We have introduced a modelling framework that provides real-time forecasts during an emerging animal health crisis, and it is designed to be rapidly adaptable to any outbreak context. Our results highlight several important considerations when using our model to guide decisions. *Forecasts made before the peak of an outbreak should be used with caution*, particularly early in the outbreak when reported caseloads are low. What can also be done at this stage is to estimate the probability with which a specific outcome may occur (e.g., the proportion of predicted trajectories that surpass a specific number of cases by a certain date, Wilke and Bergstrom 2020). *Greater emphasis should be placed on short-term forecasts of up to two weeks ahead*; as longer-term forecasts, such as peak size and final size are more difficult, particularly for early timepoints. Finally, *the ability to generate reliable and timely predictions ultimately hinges on data availability and quality*. While we have not explicitly investigated this here, common data issues ranging from reporting delays (e.g., Reich 2016, Moss et al. 2023) to unreported/undetected cases (e.g., Probert 2018) are known to diminish forecast performance.

The modelling framework and initial findings presented here opens doors for follow-up activities, which should contribute to improving animal disease forecasting capabilities and building preparedness in model deployment during future crises. Extending the framework to incorporate airborne- and vector-mediated spread (e.g., Jewell and Brown 2015) will allow application to animal disease such as FMD, LSD and HPAI, which are of high concern. Use of our framework for real-time evaluation of control strategies to guide choice of intervention can be explored (e.g., Probert 2018), noting that it already possesses the capacity to do so for vaccination and culling measures (and can be easily extended to other intervention types). In terms of improving forecast performance, numerous refinements to the epidemic model and ABC-SMC fitting algorithm can be explored, including: the addition of an observation model to account for incomplete data (e.g., undetected infections, reporting delays), integration of movement/contact network data, a spatiotemporal distance measure that compares the observed and simulated spatial distribution of infections at multiple time intervals (e.g., Minter and Retkute 2019). In addition, investigating the importance of data accuracy and completeness to forecast reliability is also a critical area of further research. Finally, as recognised in public health, there is a similar need for ongoing collaborations between modellers and animal health practitioners to fully leverage modelling tools to support outbreak response. Simulation (or synthetic) exercises held in collaboration with practitioners can provide valuable experience in real-time deployment of our modelling framework, adaptation to unexpected situations, working with incomplete and evolving data, and effective communication of forecast outputs to decision-makers. Several activities outlined here are underway through ongoing partnerships with Australian government and industry institutions, aimed at strengthening the country’s capacity for rapid decision-support during animal disease outbreaks.

## 5 Data accessibility

The C++ code implementing the simulation and R code implementing the MCMC algorithms used in this study may be found at https://gitlab.unimelb.edu.au/mtheng/livestock-forecasting, released under the GNU Public Licence v. 3. Access to the underlying data is protected to preserve the confidentiality of premises locations. Requests for these data should be directed to the NSW Department of Primary Industries.

## 6 Acknowledgements

This research was funded by the Australian Government Department of Agriculture, Fisheries and Forestry under Biosecurity2030 Project C09530: Modelling for real-time decision support in animal disease outbreaks and through the Australian Research Council’s Discovery Projects funding scheme (project DP210103239). Simin Lee was supported by the Warren Clark Bequest Veterinary Science Research Scholarship from the University of Melbourne. This research was also supported by The University of Melbourne’s Research Computing Services and the Petascale Campus Initiative. We thank Rob Moss for his advice on disease forecasting and evaluation. We also thank Thao P. Le for their feedback on the manuscript.

