## Supplementary Material for "A real-time forecasting framework for emerging infectious diseases affecting animal populations"

**References** 40

### A ABC-SMC setup

#### A.1 Simulation process

The general simulation process is as follows:

1. At the start of each simulated day of the epidemic, the states of all premises are updated to reflect the current knowledge of the system.
2. Values for unknown parameters are sampled from their respective priors.
3. The total amount of infection pressure in the system, $\tau_t$, is calculated.
4. A random sample is drawn from a Poisson distribution with mean $\tau_t$ to calculate the number of new infections proposed to occur based on our expectation $E_t^* \sim \text{Pois}(\tau_t)$.

5. Proposed new infectees ($E_t^*$) are sampled from the set of those premises susceptible at that time, $S_t$, probabilistically weighted based on the infection pressure being exerted on them by all the infectious premises at that time, $I_t$.
6. The proposed infectors ($I_t^*$) of these new infectees are sampled from $I_t$ probabilistically weighted based on the infection pressure that they are each exerting on each of the proposed infectees at time $t$.
7. The Poisson process is locally-thinned using a retrospective sampling technique that enables valid simulation of infection times across space and time (Karr, 1991). This involves only allowing each new infection to occur if the prevalence of infection on the proposed infector ($i \in I_t^*$) is greater than a random draw, $u$, from a uniform distribution $u \sim \mathcal{U}(0,1)$, i.e., if $h(t)_[i] > u$, then infection occurs at $j \in E_t^*$.
8. The states of all premises are updated, time is advanced a day ($t \to t+1$) and the process repeated from step 2.

#### A.2 ABC-SMC algorithm

Pseudo-code for our ABC-SMC algorithm are as follows (adapted from Minter and Retkute 2019):

1. Set the number of generations $G$, and the number of particles $N_{\text{px}}$.
2. Set the temporal and spatial tolerance values for each generation (temporal: $\epsilon_{\text{temp},1} > \epsilon_{\text{temp},2} > ..., > \epsilon_{\text{temp},G}$; spatial: $\epsilon_{\text{spat},1} < \epsilon_{\text{spat},2} < ..., < \epsilon_{\text{spat},G}$).
3. Set the generation indicator $g = 1$ and particle indicator $i = 1$.
4. If $g = 1$, sample $\theta^{**}$ from the prior distribution $P(\theta)$. If $g > 1$, sample $\theta^*$ from the previous generation $\theta_{g-1}$ with weights $w_{g-1}$, and perturb the particle to obtain $\theta^{**} \sim \mathcal{N}(\theta^*, \sqrt{2\tau_{g-1}^2})$.
5. Generate dataset $D^{**}$ from the model using $\theta^{**}$ and calculate the temporal ($d_{\text{temp}}(D, D^{**})$) and spatial ($d_{\text{spat}}(D, D^{**})$) distances.
6. If $d_{\text{temp}}(D, D^{**}) \leq \epsilon_{\text{temp},g}$ and $d_{\text{spat}}(D, D^{**}) \geq \epsilon_{\text{spat},g}$, accept $\theta^{**}$. Otherwise, reject and return to step 4.
7. Set $\theta_g^{(i)} = \theta^{**}$ and calculate the corresponding weight of the accepted particle $i$,

$$w_g^{(i)} = \begin{cases} \hat{P}(D|D^{**})P(\theta^{**}), & \text{if } g = 1. \\ \dfrac{\hat{P}(D|D^{**})P(\theta^{**})}{\sum_{i=1}^{N_{\text{px}}} w_{g-1} \cdot f_{\mathcal{N}}(\frac{\theta_g^{(i)} - \theta_{g-1}}{\sqrt{2\tau_{g-1}^2}} | \mu=0, \sigma=1)}, & \text{if } g > 1. \end{cases}$$

8. If $i < N_{\text{px}}$, make the increment $i = i + 1$ and go to step 4.
9. Normalise the weights so that $\sum_{k=1}^{N_{\text{px}}} w_g^{(i)} = 1$.
10. If $g < G$, set $g = g + 1$, go to step 3.

Note that the distance functions, sequence of tolerance values, number of generations, number of simulations for each parameter set, and perturbation kernel to use are user-defined choices that are context dependent (see Minter and Retkute 2019). These choices not only determine how efficiently the parameter space is explored, it can also affect the model's goodness-of-fit to the data.

Table A1: Parameter values for (i) the transmission model and (ii) the ABC-SMC.

| | **Parameter(s)** | **Description** | **Value** |
|---|---|---|---|
| (i) | $t_{\text{detect}}$ | Day of first detection/notification (indexed) | 9, 10, 10, 12[a] |
| | $t_{\text{today}}$ | Data date/time point | 21, 35, 49 days after $t_{\text{detect}}$ |
| | $t_{\text{lag}}$ | Number of days before the current data date to fit model up to | 3 |
| | $\beta_{\text{intra}}$ | Within-premises transmission rate | $\sim \mathcal{U}(1, 20)$ |
| | $\sigma_{\text{intra}}$ | Within-premises latency rate = 1/latent period | 1/1 |
| | $\gamma_{\text{intra}}$ | Within-premises recovery rate = 1/infectious period | 1/6 |
| | $\alpha$ | Background transmission rate | $\sim \mathcal{U}(0, 0.02)$ |
| | $\beta_0$ | Baseline transmission rate (from S to E at premises-level) | $\sim \mathcal{U}(0, 0.02)$ |
| | $\psi$ | Kernel shape parameter (decay of # with distance between premises) | $\sim \mathcal{U}(0, 20)$ |
| | $\zeta$ | Index of the count or density of animals on premises i at time t | $\sim \mathcal{U}(-2, 2)$ |
| | $\xi$ | Index of the count or density of animals on premises i at time t | $\sim \mathcal{U}(-2, 2)$ |
| | $\theta$ | Effectiveness of vaccination | $\sim \mathcal{U}(0, 1)$ |
| (ii) | $N_{\text{px}}$ | Number of accepted particles | 1,000 |
| | $G$ | Maximum number of ABC-SMC generations/steps | 8 |
| | $\epsilon_{\text{temp},1}, \epsilon_{\text{temp},2}, ..., \epsilon_{\text{temp},8}$ | Temporal tolerance schedule (+/- daily case counts) | 50, 40, 30, 20, 15, 10, 7.5, 6 |

| Parameter(s) | Description | Value |
|---|---|---|
| $\epsilon_{\text{spat},1}, \epsilon_{\text{spat},2}, \ldots, \epsilon_{\text{spat},8}$ | Spatial tolerance schedule (Pearson's correlation of grid cell case counts between observed and simulated data) | 0.3, 0.4, 0.5, 0.6, 0.65, 0.675, 0.7, 0.725 |
| $X \times X$ | Number of grids (for grid cell case counts) | $20 \times 20$ |

[a] Day 1 being the estimated date of first infection

### B Extended results

#### B.1 Forecast targets

Forecasts of summary statistics by cluster and time point are given in Table B1.

#### B.2 Additional runs

We produced forecasts for the Central Coast-Maitland-Newcastle (CCMN) cluster at two additional timepoints beyond the observed peak (9- and 11-weeks in). Figure B1 shows that though these forecasts no longer predicted a much larger outbreak, the model struggled to fit the peak and predicted a relatively smaller but protracted outbreak.

#### B.3 Spatial forecasts

Spatial forecasts for all clusters/regions and timepoints can be found in following Figures B2–B17. Note that we have also included spatial predictions for the full dataset (i.e., $t_{\text{today}} = t_{\max} - 1$) for each cluster as a reference for the "best" fit the model can achieve.

Table B1: Predictions of summary statistics by cluster. Median estimates accompanied by their 95% credible intervals (in brackets) are provided. Summary statistics predicted were: final outbreak size ('final_size'), timing of peak ('peak_t'), case count at peak ('peak_n'), and outbreak duration ('duration').

(a) Sydney Basin

| | Predicted (3 weeks) | Predicted (5 weeks) | Predicted (7 weeks) | Observed |
|---|---|---|---|---|
| final_size | 6702 (4162,7350) | 3568 (2795,6243) | 3054 (2649,3467) | 2875 |
| peak_t | 42 (36,49) | 37 (24,57) | 31 (24,39) | 26 |
| peak_n | 230 (121,382) | 98 (71,132) | 96 (72,122) | 98 |
| duration | 125 (83,188) | 154 (117,196) | 135 (103,185) | 126 |

(b) Central Coast-Maitland-Newcastle

| | Predicted (3 weeks) | Predicted (5 weeks) | Predicted (7 weeks) | Observed |
|---|---|---|---|---|
| final_size | 948 (9,6548) | 5268 (3929,6157) | 5160 (3628,6052) | 1292 |
| peak_t | 40 (7,166) | 71 (59,92) | 78 (63,97) | 53 |
| peak_n | 57 (3,6471) | 132 (75,219) | 100 (55,160) | 48 |
| duration | 77 (12,185) | 176 (134,185) | 184 (165,185) | 128 |

(c) Tamworth

| | Predicted (3 weeks) | Predicted (5 weeks) | Predicted (7 weeks) | Observed |
|---|---|---|---|---|
| final_size | 1366 (61,2317) | 1202 (652,2013) | 912 (524,1598) | 582 |
| peak_t | 58 (12,130) | 52 (30,76) | 41 (24,94) | 31 |
| peak_n | 34 (5,768) | 30 (21,57) | 22 (16,34) | 29 |
| duration | 153 (45,202) | 160 (106,202) | 178 (101,202) | 106 |

(d) Hunter Valley

| | Predicted (3 weeks) | Predicted (5 weeks) | Predicted (7 weeks) | Observed |
|---|---|---|---|---|
| final_size | 243 (100,1061) | 276 (212,365) | 306 (230,794) | 404 |
| peak_t | 35 (11,76) | 31 (24,45) | 36 (23,63) | 31 |
| peak_n | 11 (5,70) | 14 (10,19) | 15 (9,23) | 21 |
| duration | 99 (66,189) | 91 (69,138) | 98 (73,188) | 105 |

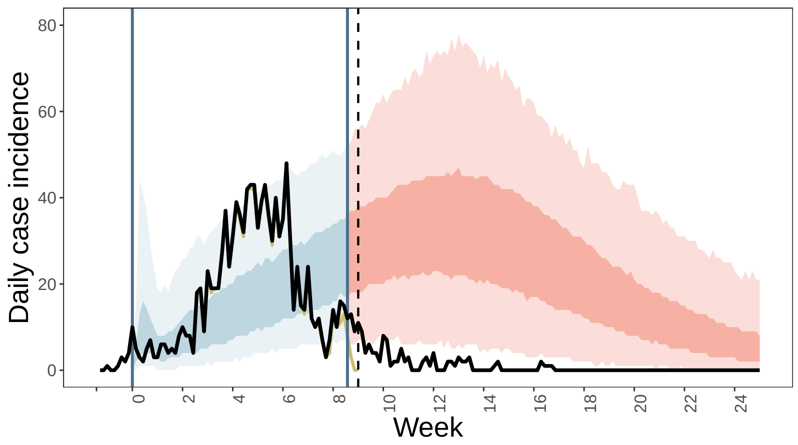

(a)

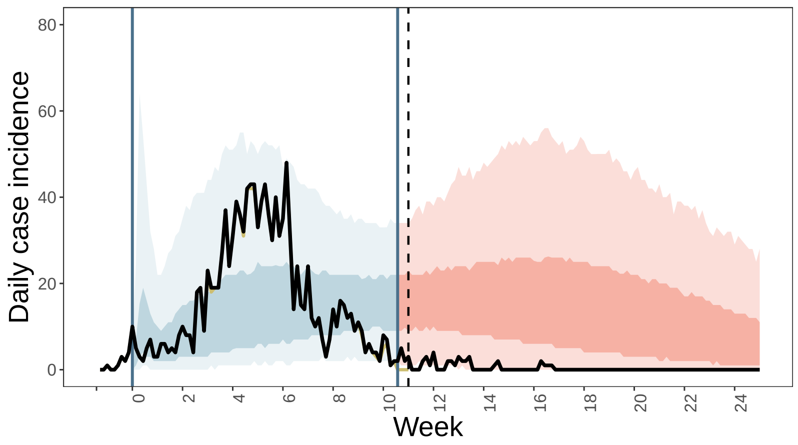

(b)

Figure B1: Forecast of daily EI case incidence for the Central Coast-Maitland-Newcastle (CCMN) cluster at two additional timepoints beyond the observed peak: (a) 9-weeks in, and (b) 11-weeks in. The blue shaded regions represent model fit to the data available at the time of forecasting (dark yellow lines). The red shaded regions represent the forecast. Vertical black dashed lines indicate the time point. Vertical blue lines indicate the time window of data the model was fit to (accounting for delay with a 3-day lag). Black lines represent all reported daily case counts (i.e., all observations by end of outbreak). Shaded areas indicate the 50% and 95% percentile of 1000 simulated trajectories (of accepted particles).

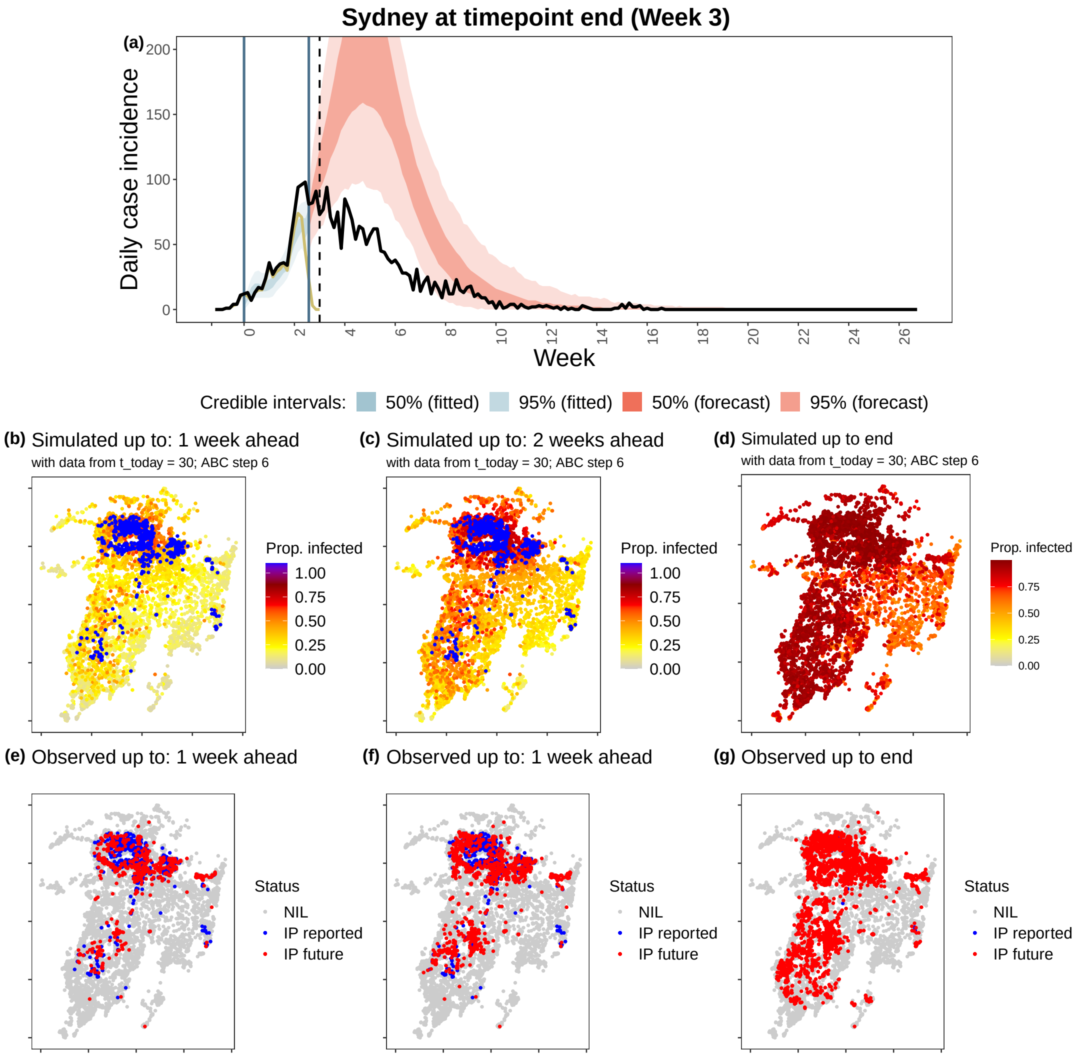

Figure B2: Comparison of simulated and observed outbreaks for the **Sydney Basin region at timepoint 1 (week 3)**. (a) Simulated epidemic curve, with 50% and 95% credible interval (shaded regions), compared to all observations (black line) and to only observations available at the time of forecasting (dark yellow line). Spatial maps showing proportion of runs (out of 1000 simulation runs) in which a premises was infected between the date of first detection and (b) 1-week ahead from the current date, (c) 2-weeks ahead from the current date, and (d) the end of the outbreak; compared to (e-g) the single observed realisation of the outbreak at the same respective dates ("NIL": non-infected premises; "IP reported": infected premises reported before the timepoint; "IP future": infected premises reported after the timepoint).

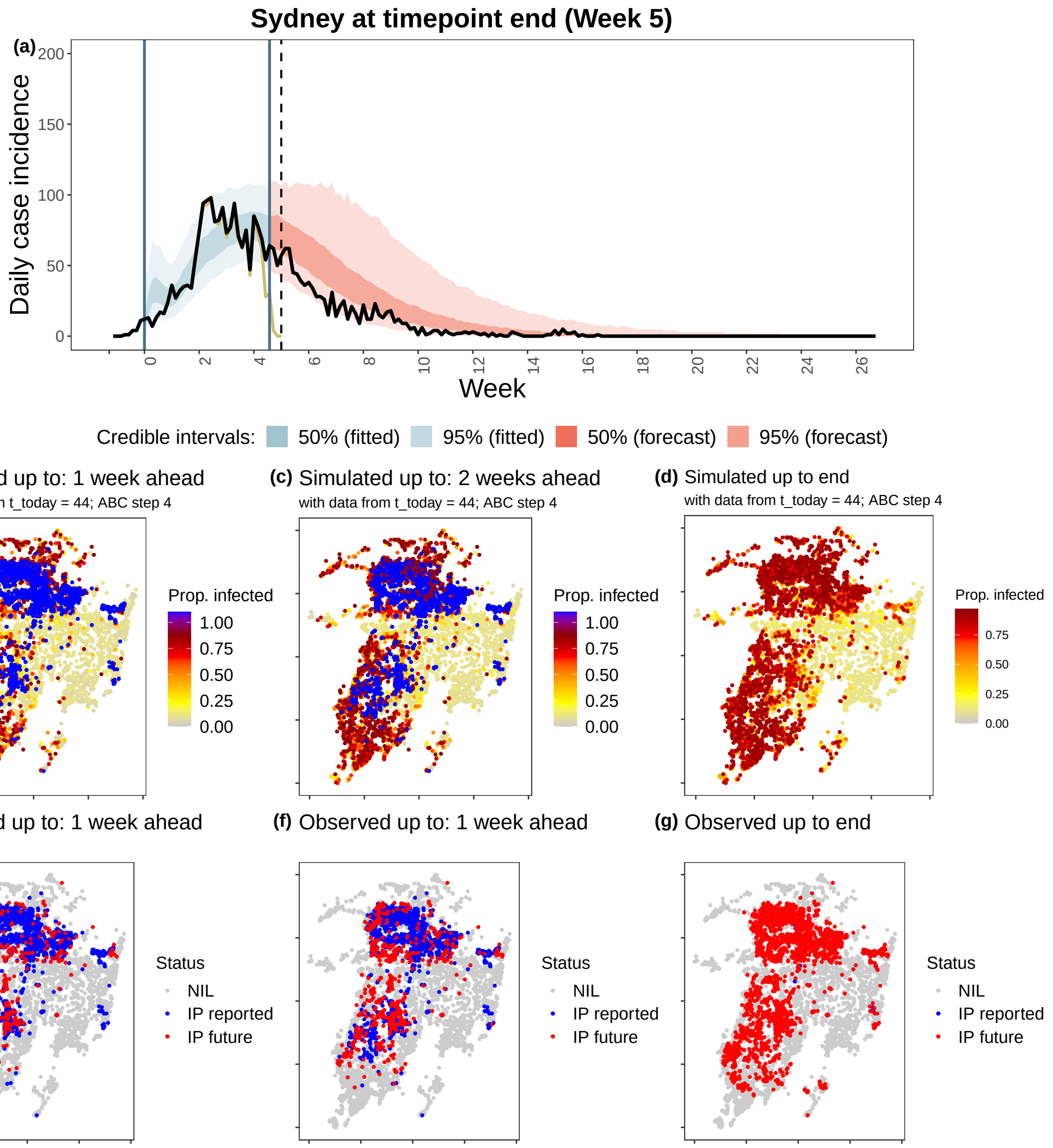

Figure B3: Comparison of simulated and observed outbreaks for the **Sydney Basin region at timepoint 2 (week 5)**. (a) Simulated epidemic curve, with 50% and 95% credible interval (shaded regions), compared to all observations (black line) and to only observations available at the time of forecasting (dark yellow line). Spatial maps showing proportion of runs (out of 1000 simulation runs) in which a premises was infected between the date of first detection and (b) 1-week ahead from the current date, (c) 2-weeks ahead from the current date, and (d) the end of the outbreak; compared to (e-g) the single observed realisation of the outbreak at the same respective dates ("NIL": non-infected premises; "IP reported": infected premises reported before the timepoint; "IP future": infected premises reported after the timepoint).

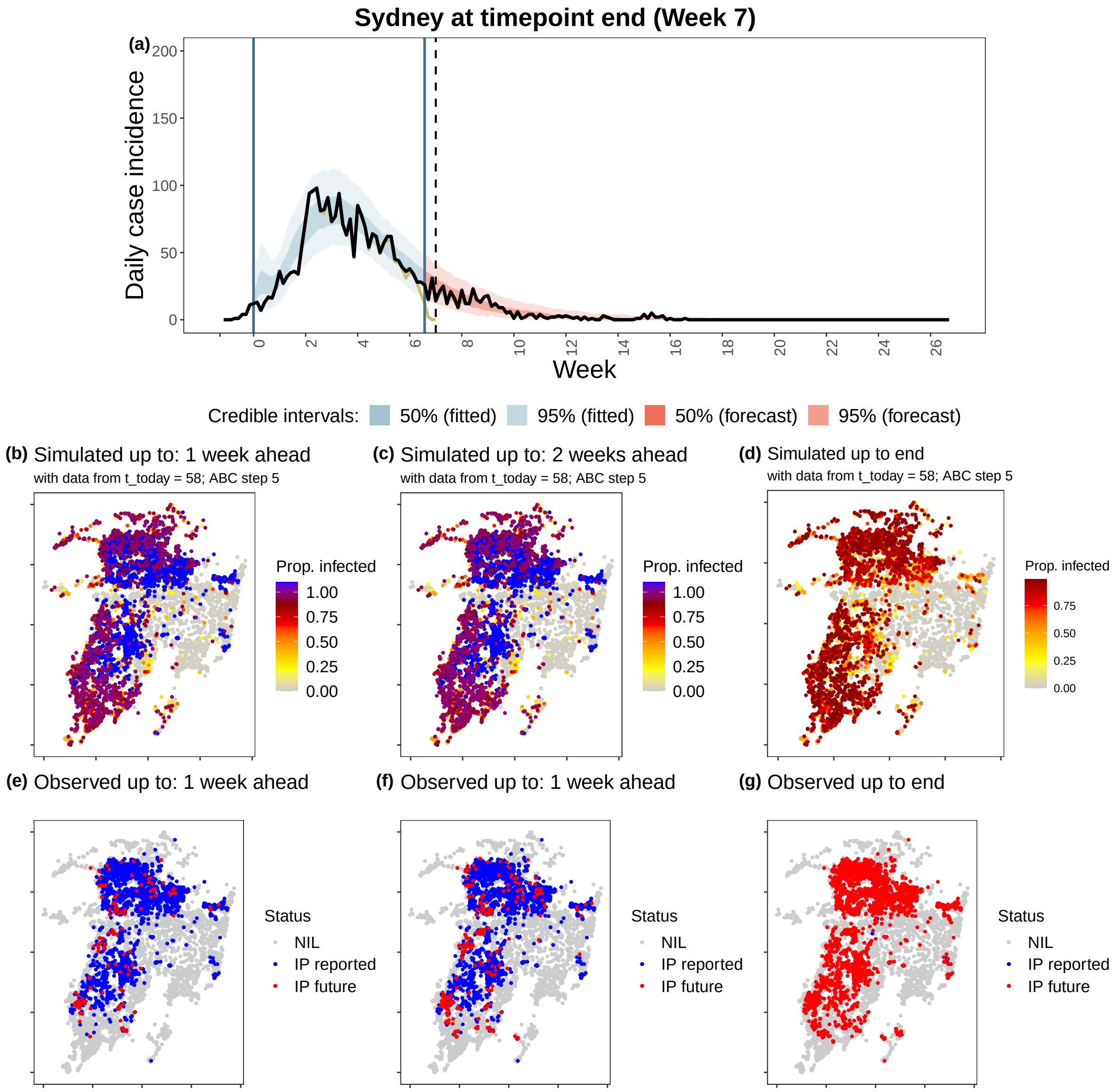

Figure B4: Comparison of simulated and observed outbreaks for the **Sydney Basin region at timepoint 3 (week 7)**. (a) Simulated epidemic curve, with 50% and 95% credible interval (shaded regions), compared to all observations (black line) and to only observations available at the time of forecasting (dark yellow line). Spatial maps showing proportion of runs (out of 1000 simulation runs) in which a premises was infected between the date of first detection and (b) 1-week ahead from the current date, (c) 2-weeks ahead from the current date, and (d) the end of the outbreak; compared to (e-g) the single observed realisation of the outbreak at the same respective dates ("NIL": non-infected premises; "IP reported": infected premises reported before the timepoint; "IP future": infected premises reported after the timepoint).

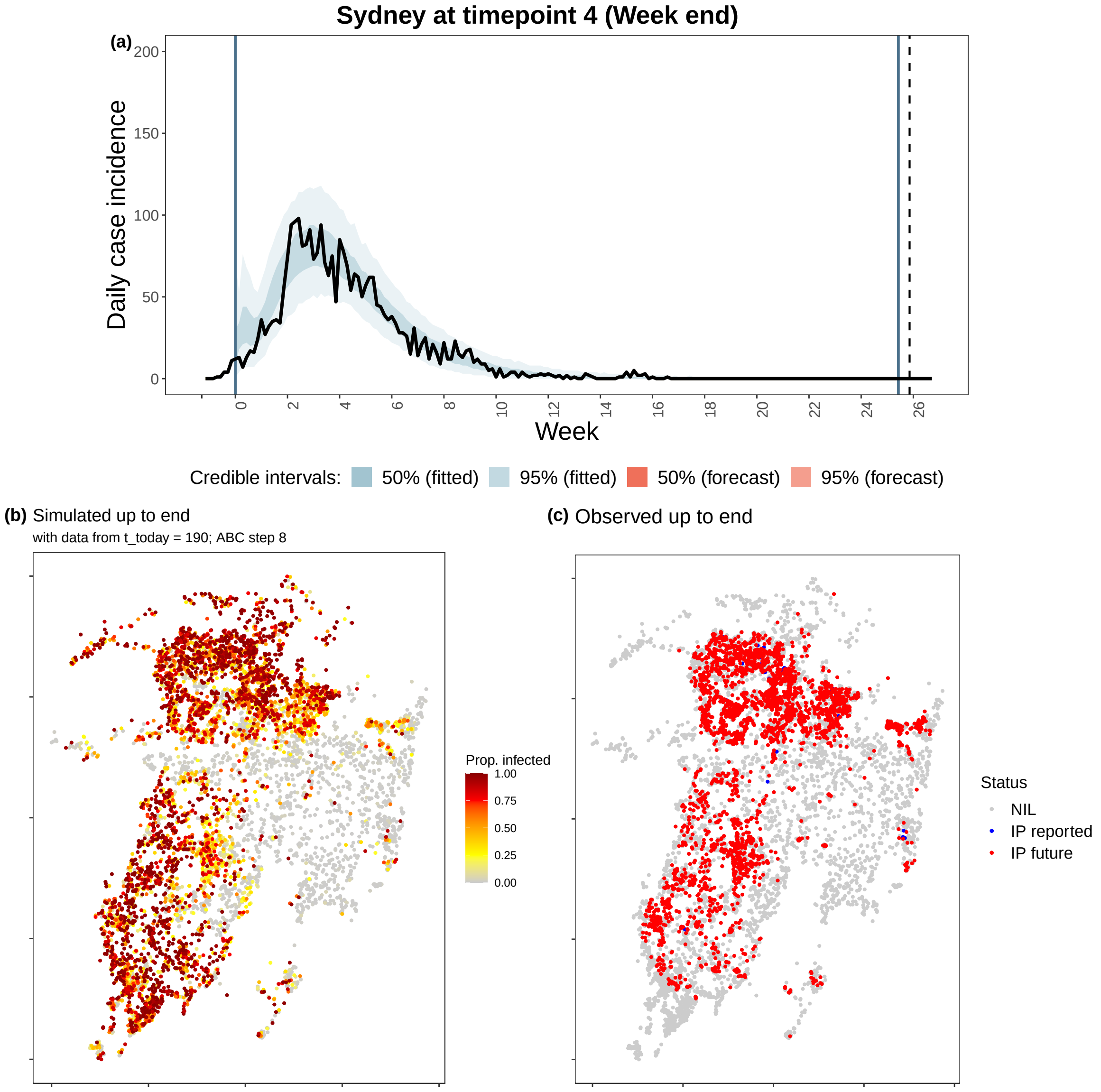

Figure B5: Comparison of simulated and observed outbreaks for the **Sydney Basin region at timepoint 4 (end)**. (a) Simulated epidemic curve, with 50% and 95% credible interval (shaded regions), compared to all observations (black line) and to only observations available at the time of forecasting (dark yellow line). Spatial maps showing proportion of runs (out of 1000 simulation runs) in which a premises was infected between the date of first detection and (b) the end of the outbreak, (c) the single observed realisation of the outbreak at the end ("NIL": non-infected premises; "IP reported": infected premises reported before the timepoint; "IP future": infected premises reported after the timepoint).

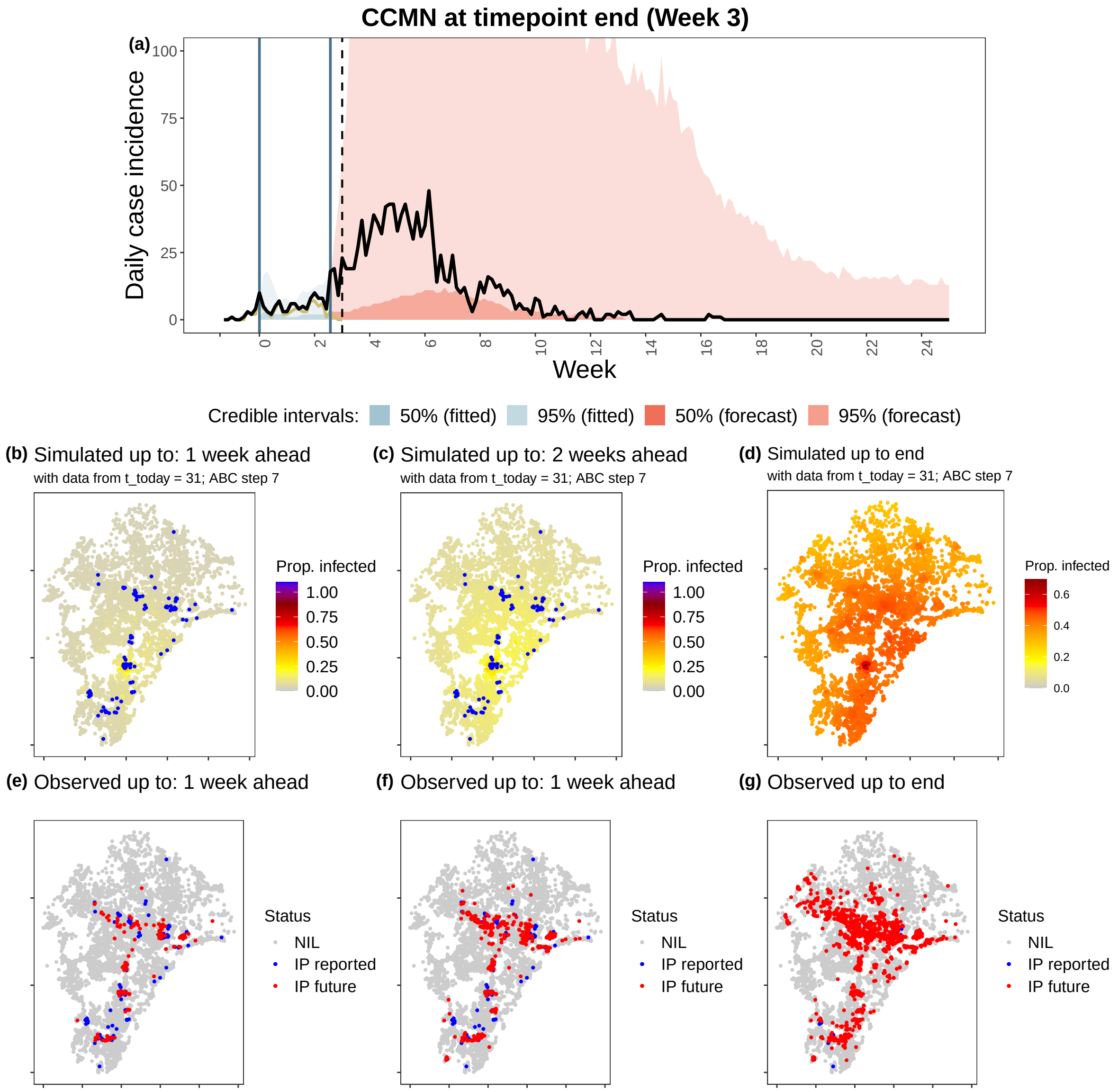

Figure B6: Comparison of simulated and observed outbreaks for the **Central Coast-Maitland-Newcastle region at timepoint 1 (week 3)**. (a) Simulated epidemic curve, with 50% and 95% credible interval (shaded regions), compared to all observations (black line) and to only observations available at the time of forecasting (dark yellow line). Spatial maps showing proportion of runs (out of 1000 simulation runs) in which a premises was infected between the date of first detection and (b) 1-week ahead from the current date, (c) 2-weeks ahead from the current date, and (d) the end of the outbreak; compared to (e-g) the single observed realisation of the outbreak at the same respective dates ("NIL": non-infected premises; "IP reported": infected premises reported before the timepoint; "IP future": infected premises reported after the timepoint).

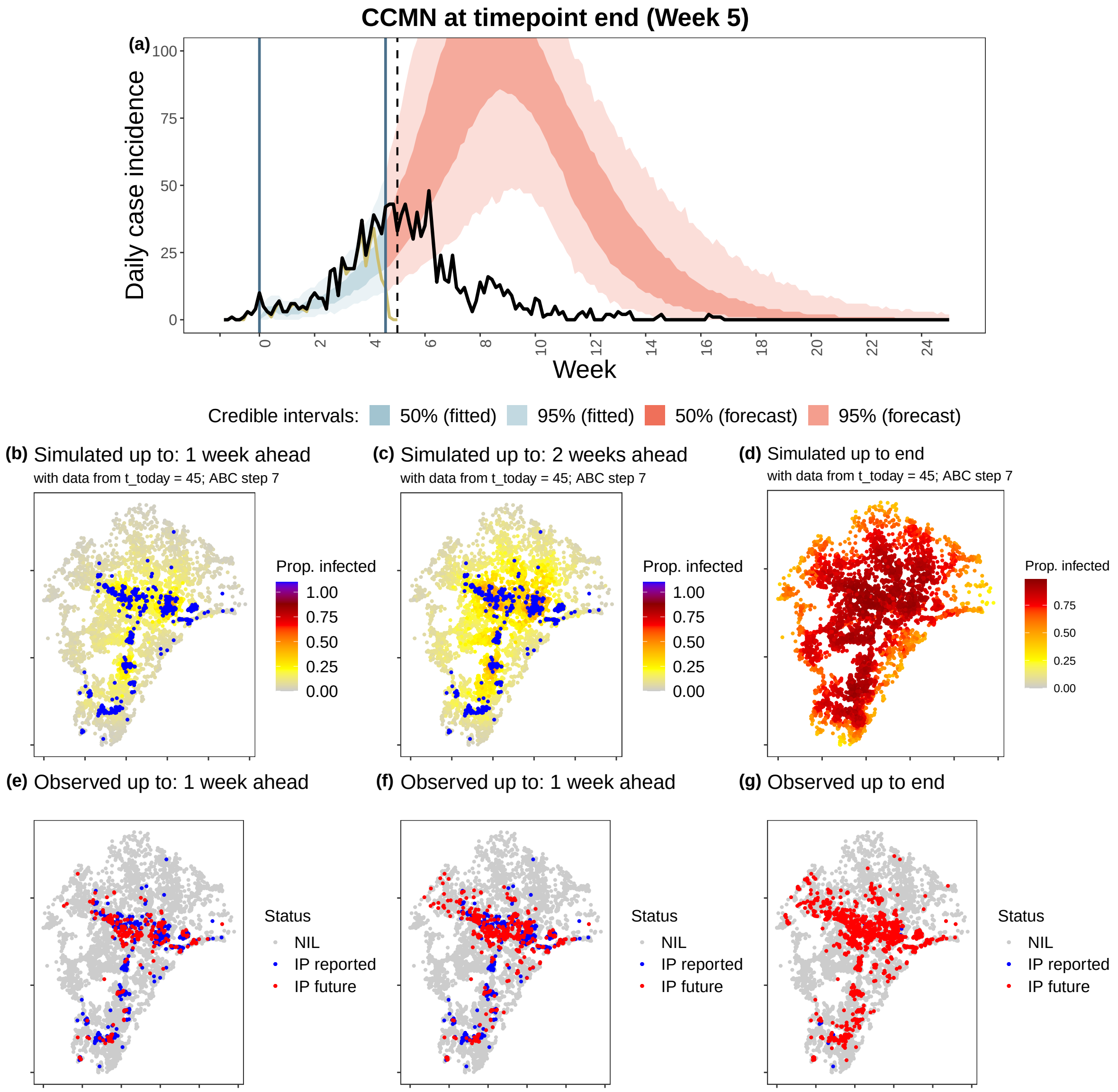

Figure B7: Comparison of simulated and observed outbreaks for the **Central Coast-Maitland-Newcastle region at timepoint 2 (week 5)**. (a) Simulated epidemic curve, with 50% and 95% credible interval (shaded regions), compared to all observations (black line) and to only observations available at the time of forecasting (dark yellow line). Spatial maps showing proportion of runs (out of 1000 simulation runs) in which a premises was infected between the date of first detection and (b) 1-week ahead from the current date, (c) 2-weeks ahead from the current date, and (d) the end of the outbreak; compared to (e-g) the single observed realisation of the outbreak at the same respective dates ("NIL": non-infected premises; "IP reported": infected premises reported before the timepoint; "IP future": infected premises reported after the timepoint).

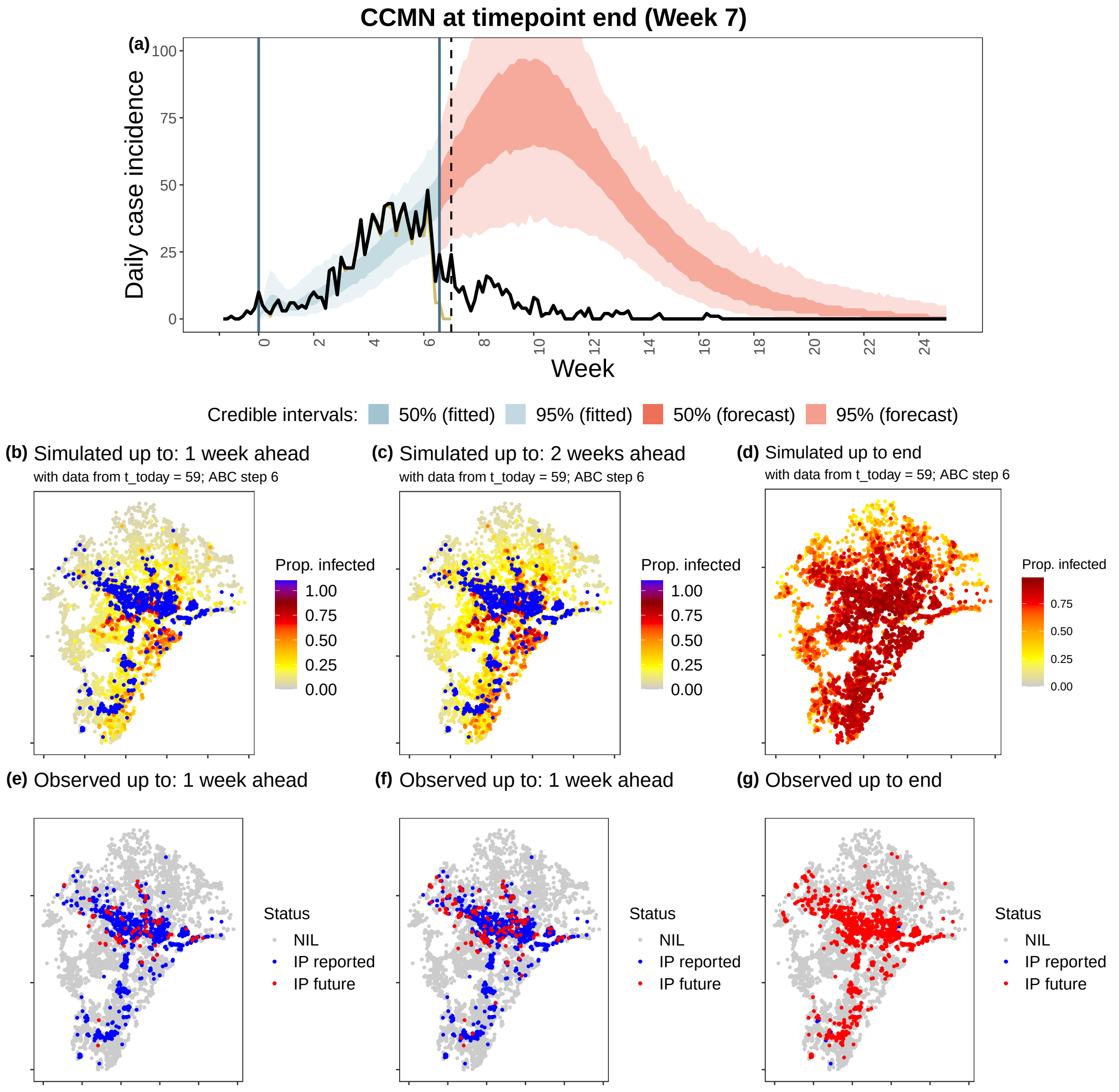

Figure B8: Comparison of simulated and observed outbreaks for the **Central Coast-Maitland-Newcastle region at timepoint 3 (week 7)**. (a) Simulated epidemic curve, with 50% and 95% credible interval (shaded regions), compared to all observations (black line) and to only observations available at the time of forecasting (dark yellow line). Spatial maps showing proportion of runs (out of 1000 simulation runs) in which a premises was infected between the date of first detection and (b) 1-week ahead from the current date, (c) 2-weeks ahead from the current date, and (d) the end of the outbreak; compared to (e-g) the single observed realisation of the outbreak at the same respective dates ("NIL": non-infected premises; "IP reported": infected premises reported before the timepoint; "IP future": infected premises reported after the timepoint).

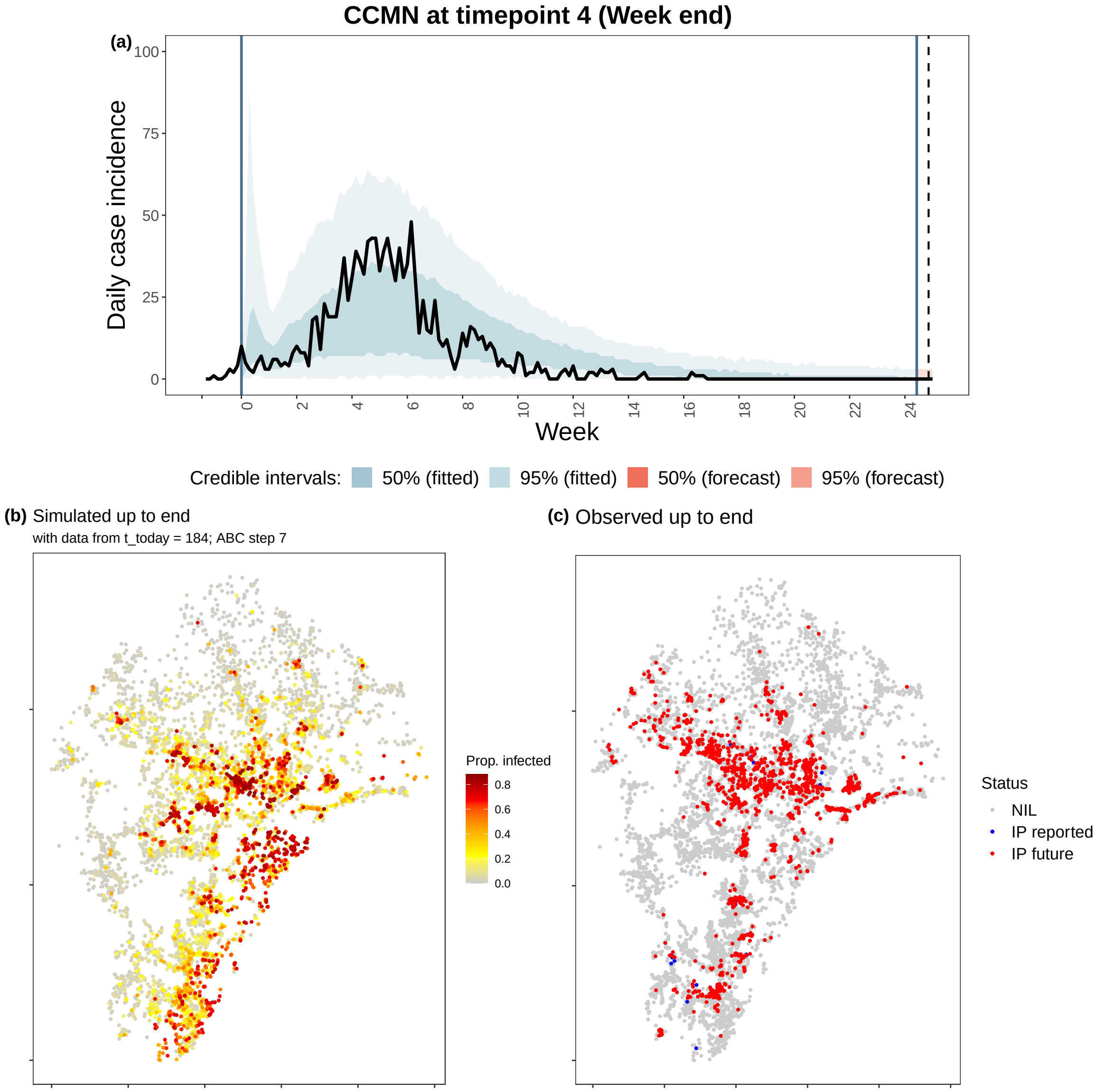

Figure B9: Comparison of simulated and observed outbreaks for the **Central Coast-Maitland-Newcastle region at timepoint 4 (end)**. (a) Simulated epidemic curve, with 50% and 95% credible interval (shaded regions), compared to all observations (black line) and to only observations available at the time of forecasting (dark yellow line). Spatial maps showing proportion of runs (out of 1000 simulation runs) in which a premises was infected between the date of first detection and (b) the end of the outbreak, (c) the single observed realisation of the outbreak at the end ("NIL": non-infected premises; "IP reported": infected premises reported before the timepoint; "IP future": infected premises reported after the timepoint).

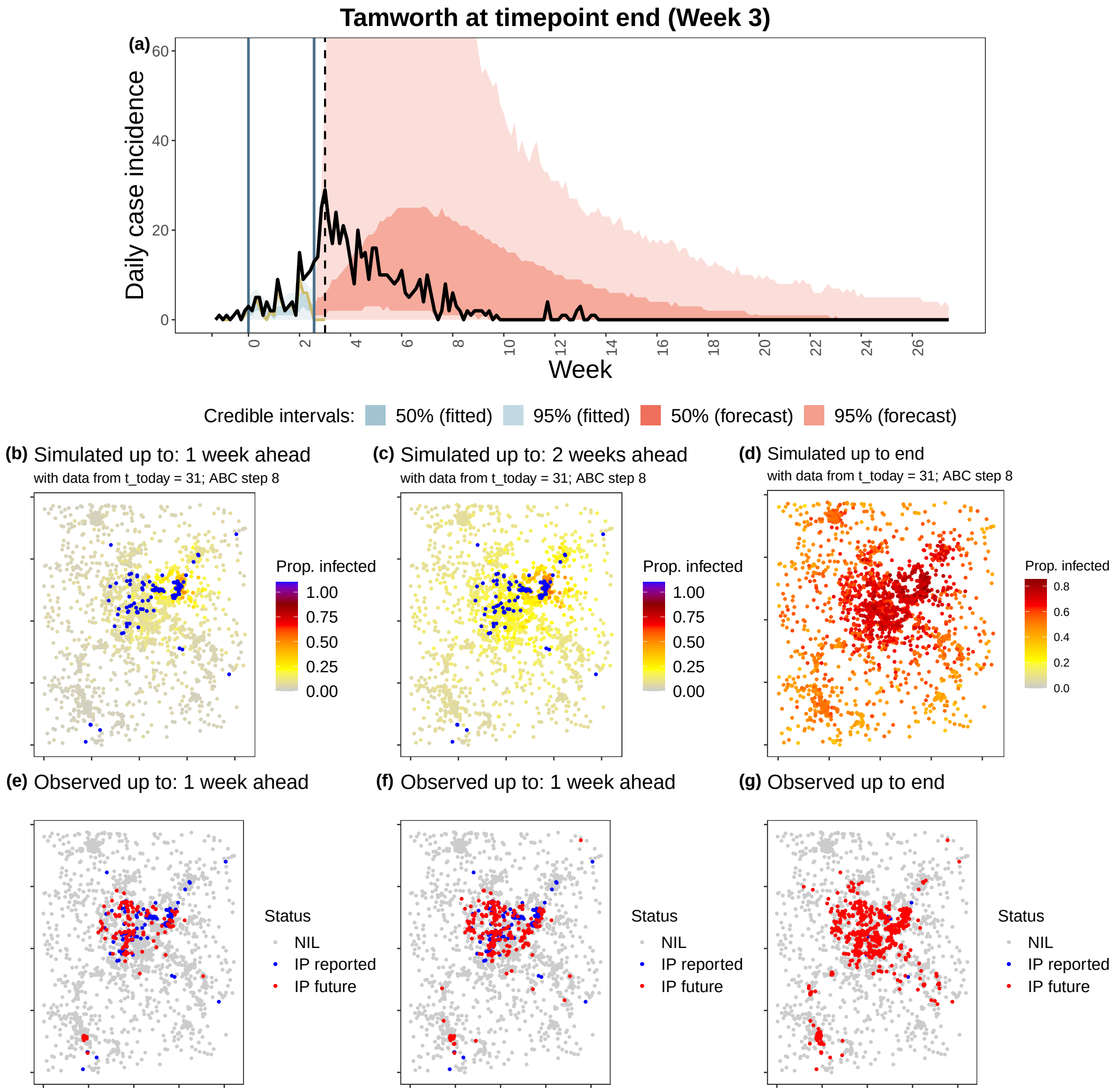

Figure B10: Comparison of simulated and observed outbreaks for the **Tamworth region at timepoint 1 (week 3)**. (a) Simulated epidemic curve, with 50% and 95% credible interval (shaded regions), compared to all observations (black line) and to only observations available at the time of forecasting (dark yellow line). Spatial maps showing proportion of runs (out of 1000 simulation runs) in which a premises was infected between the date of first detection and (b) 1-week ahead from the current date, (c) 2-weeks ahead from the current date, and (d) the end of the outbreak; compared to (e-g) the single observed realisation of the outbreak at the same respective dates ("NIL": non-infected premises; "IP reported": infected premises reported before the timepoint; "IP future": infected premises reported after the timepoint).

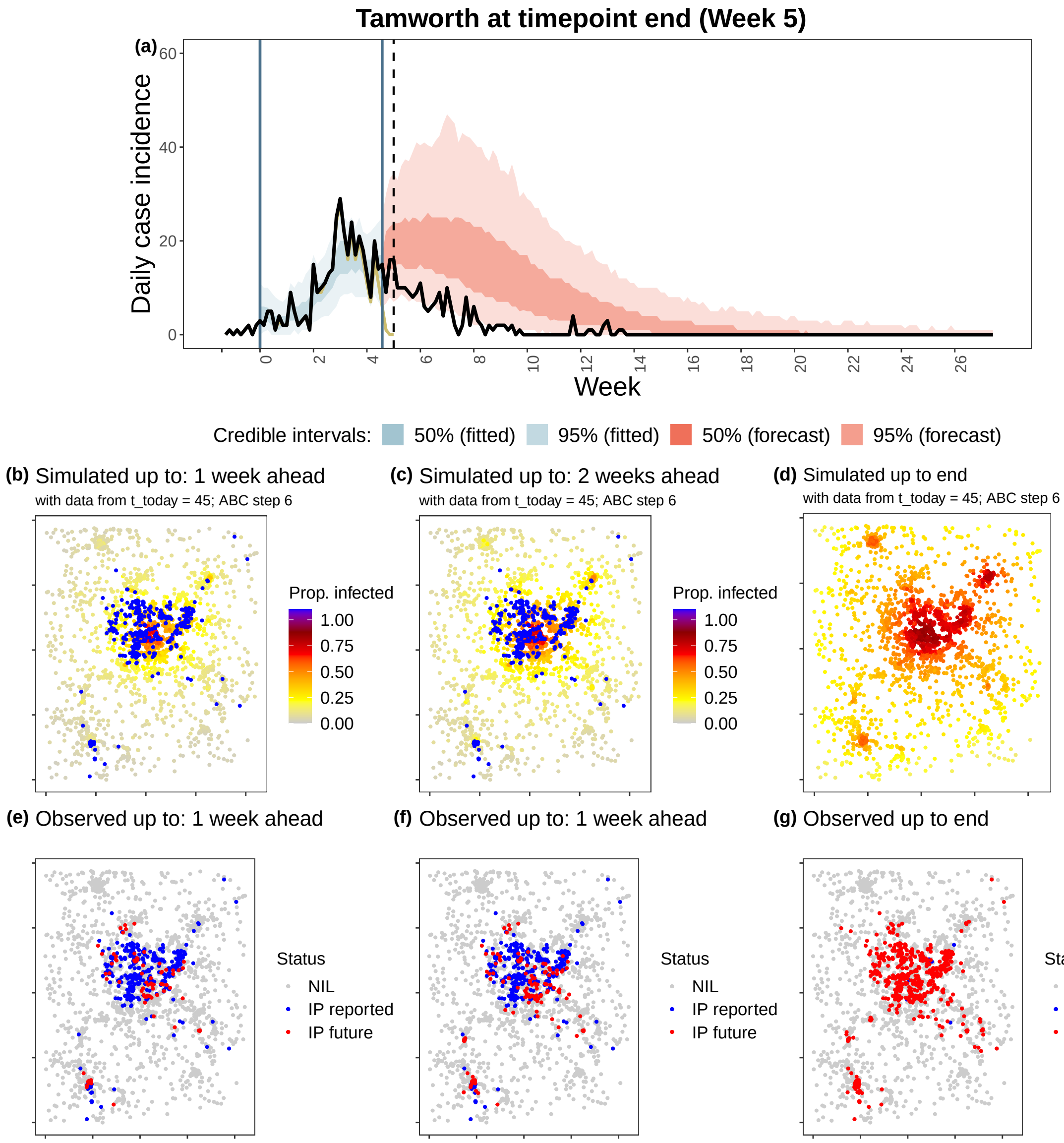

Figure B11: Comparison of simulated and observed outbreaks for the **Tamworth region at timepoint 2 (week 5)**. (a) Simulated epidemic curve, with 50% and 95% credible interval (shaded regions), compared to all observations (black line) and to only observations available at the time of forecasting (dark yellow line). Spatial maps showing proportion of runs (out of 1000 simulation runs) in which a premises was infected between the date of first detection and (b) 1-week ahead from the current date, (c) 2-weeks ahead from the current date, and (d) the end of the outbreak; compared to (e-g) the single observed realisation of the outbreak at the same respective dates ("NIL": non-infected premises; "IP reported": infected premises reported before the timepoint; "IP future": infected premises reported after the timepoint).

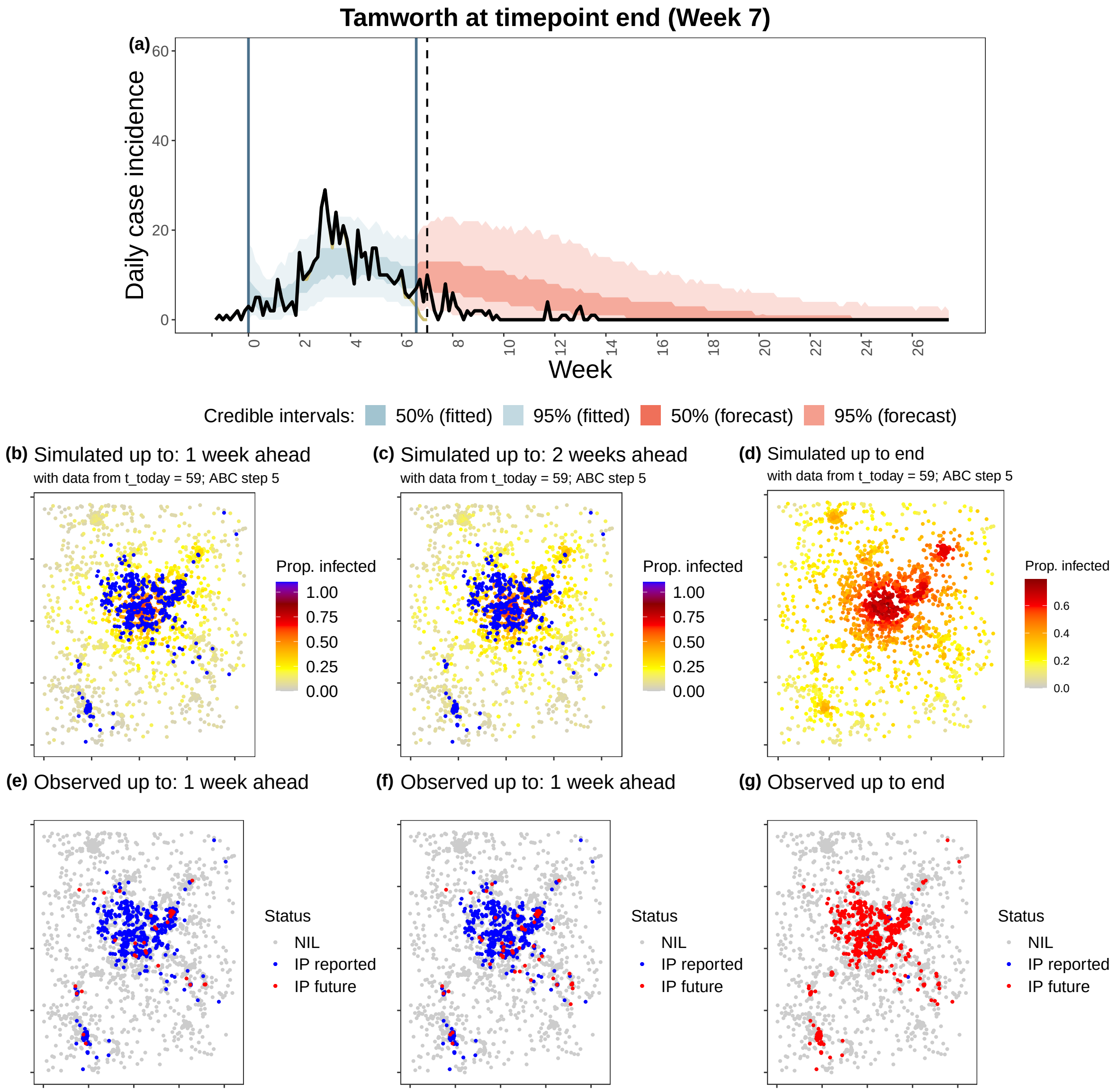

Figure B12: Comparison of simulated and observed outbreaks for the **Tamworth region at timepoint 3 (week 7)**. (a) Simulated epidemic curve, with 50% and 95% credible interval (shaded regions), compared to all observations (black line) and to only observations available at the time of forecasting (dark yellow line). Spatial maps showing proportion of runs (out of 1000 simulation runs) in which a premises was infected between the date of first detection and(b) 1-week ahead from the current date, (c) 2-weeks ahead from the current date, and (d) the end of the outbreak; compared to (e-g) the single observed realisation of the outbreak at the same respective dates ("NIL": non-infected premises; "IP reported": infected premises reported before the timepoint; "IP future": infected premises reported after the timepoint).

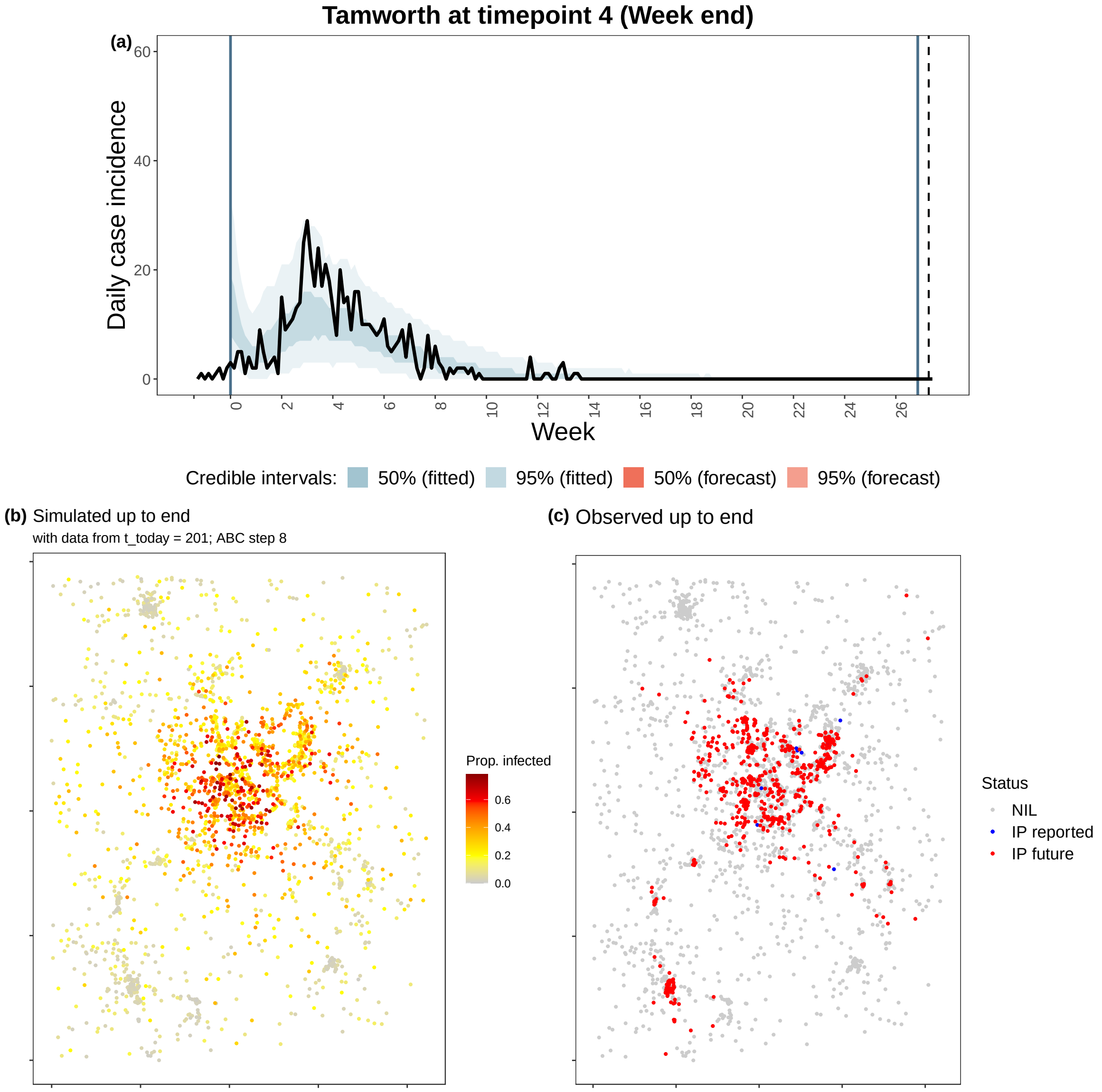

Figure B13: Comparison of simulated and observed outbreaks for the **Tamworth region at timepoint 4 (end)**. (a) Simulated epidemic curve, with 50% and 95% credible interval (shaded regions), compared to all observations (black line) and to only observations available at the time of forecasting (dark yellow line). Spatial maps showing proportion of runs (out of 1000 simulation runs) in which a premises was infected between the date of first detection and (b) the end of the outbreak, (c) the single observed realisation of the outbreak at the end ("NIL": non-infected premises; "IP reported": infected premises reported before the timepoint; "IP future": infected premises reported after the timepoint).

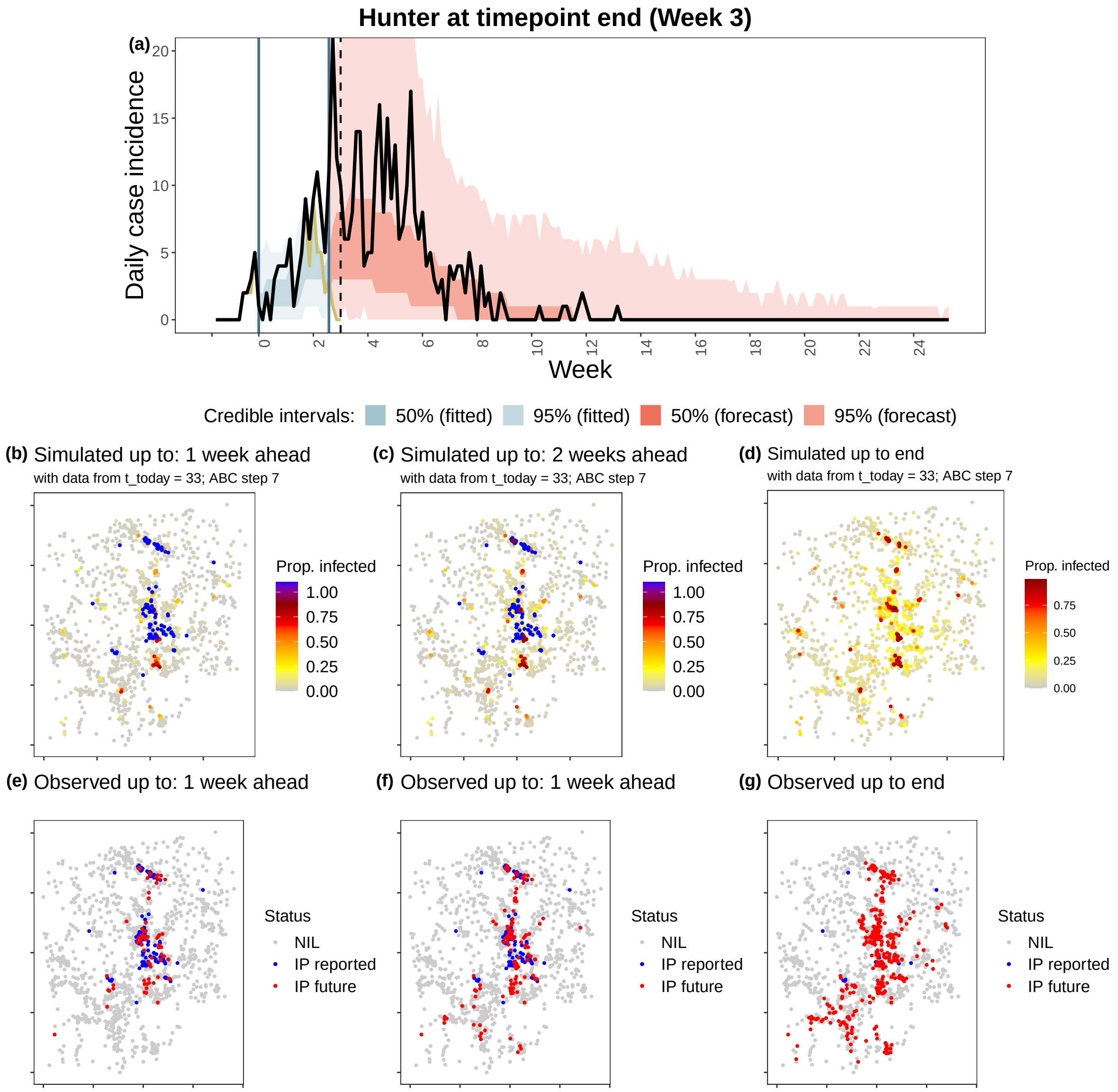

Figure B14: Comparison of simulated and observed outbreaks for the **Hunter Valley region at timepoint 1 (week 3)**. (a) Simulated epidemic curve, with 50% and 95% credible interval (shaded regions), compared to all observations (black line) and to only observations available at the time of forecasting (dark yellow line). Spatial maps showing proportion of runs (out of 1000 simulation runs) in which a premises was infected between the date of first detection and (b) 1-week ahead from the current date, (c) 2-weeks ahead from the current date, and (d) the end of the outbreak; compared to (e-g) the single observed realisation of the outbreak at the same respective dates ("NIL": non-infected premises; "IP reported": infected premises reported before the timepoint; "IP future": infected premises reported after the timepoint).

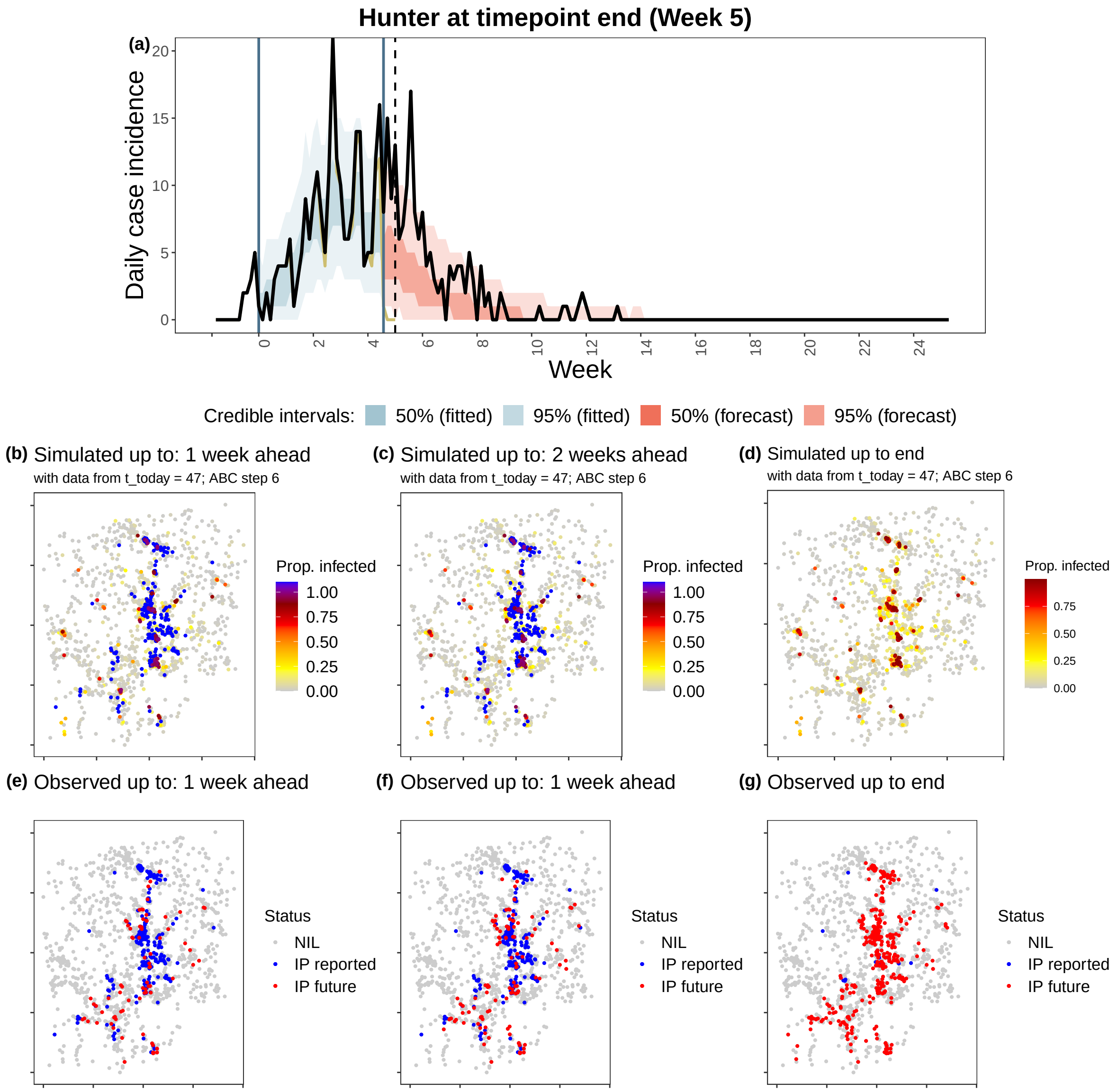

Figure B15: Comparison of simulated and observed outbreaks for the **Hunter Valley region at timepoint 2 (week 5)**. (a) Simulated epidemic curve, with 50% and 95% credible interval (shaded regions), compared to all observations (black line) and to only observations available at the time of forecasting (dark yellow line). Spatial maps showing proportion of runs (out of 1000 simulation runs) in which a premises was infected between the date of first detection and (b) 1-week ahead from the current date, (c) 2-weeks ahead from the current date, and (d) the end of the outbreak; compared to (e-g) the single observed realisation of the outbreak at the same respective dates ("NIL": non-infected premises; "IP reported": infected premises reported before the timepoint; "IP future": infected premises reported after the timepoint).

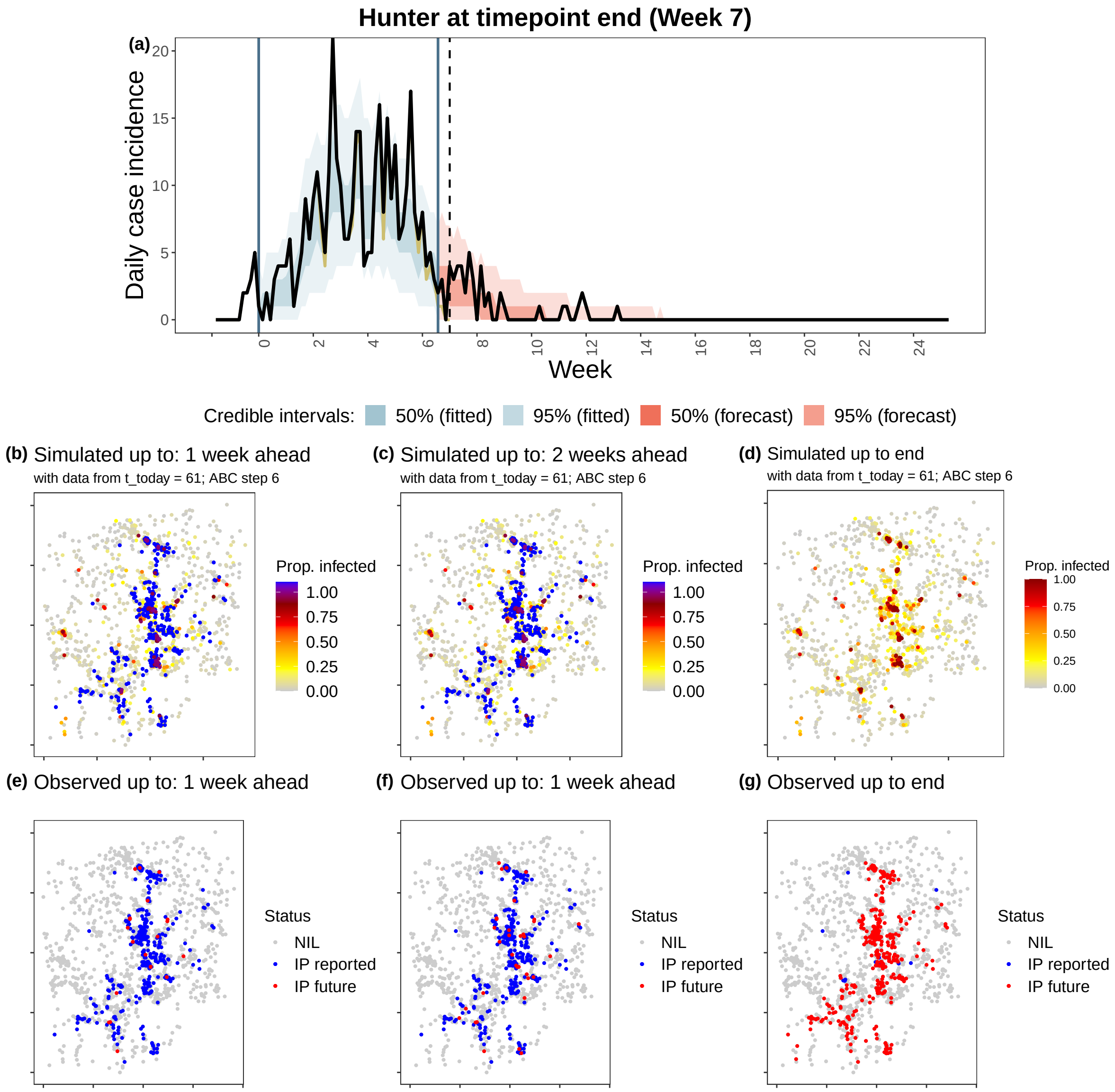

Figure B16: Comparison of simulated and observed outbreaks for the **Hunter Valley region at timepoint 3 (week 7)**. (a) Simulated epidemic curve, with 50% and 95% credible interval (shaded regions), compared to all observations (black line) and to only observations available at the time of forecasting (dark yellow line). Spatial maps showing proportion of runs (out of 1000 simulation runs) in which a premises was infected between the date of first detection and (b) 1-week ahead from the current date, (c) 2-weeks ahead from the current date, and (d) the end of the outbreak; compared to (e-g) the single observed realisation of the outbreak at the same respective dates ("NIL": non-infected premises; "IP reported": infected premises reported before the timepoint; "IP future": infected premises reported after the timepoint).

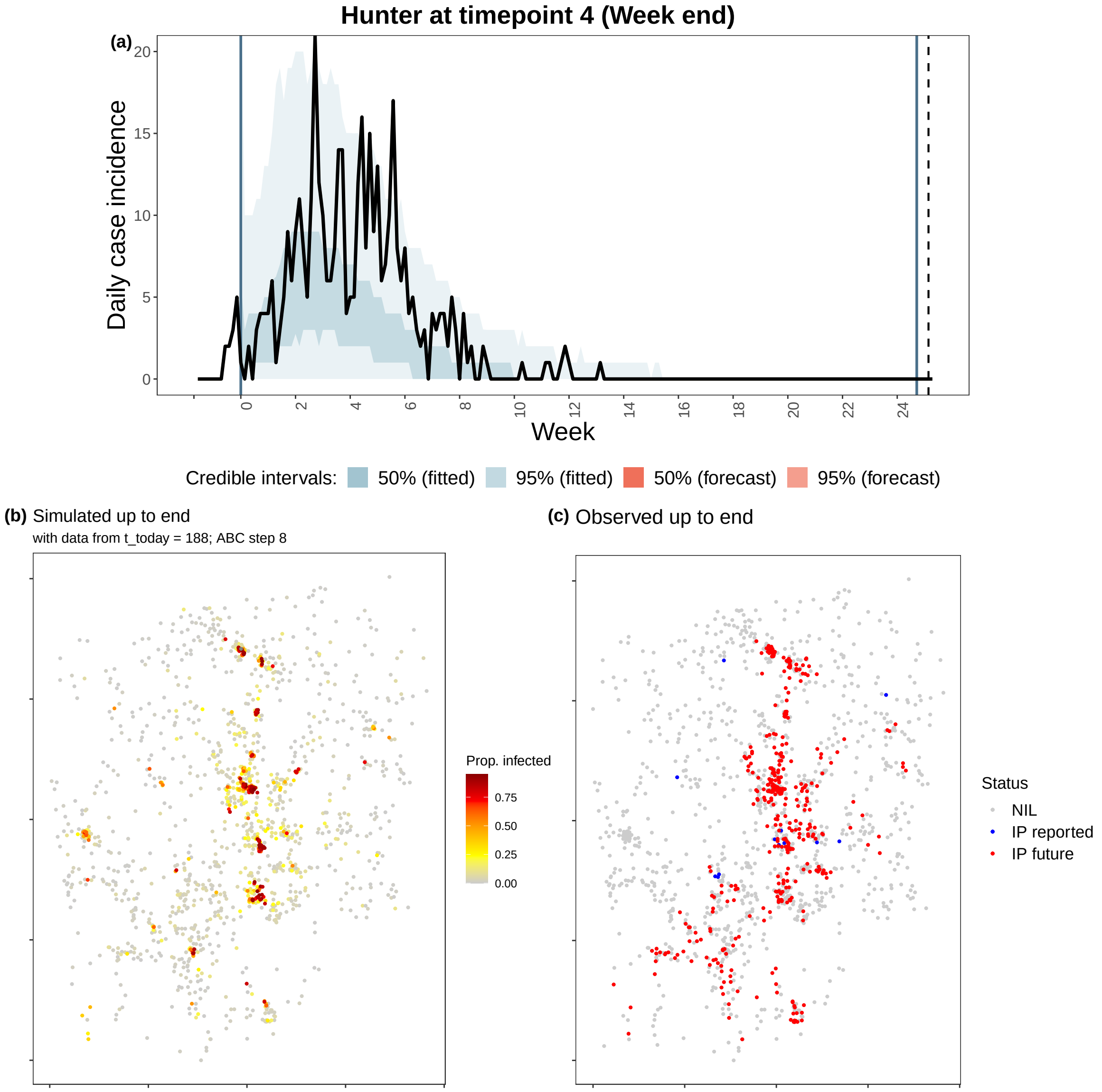

Figure B17: Comparison of simulated and observed outbreaks for the **Hunter Valley region at timepoint 4 (end)**. (a) Simulated epidemic curve, with 50% and 95% credible interval (shaded regions), compared to all observations (black line) and to only observations available at the time of forecasting (dark yellow line). Spatial maps showing proportion of runs (out of 1000 simulation runs) in which a premises was infected between the date of first detection and (b) the end of the outbreak, (c) the single observed realisation of the outbreak at the end ("NIL": non-infected premises; "IP reported": infected premises reported before the timepoint; "IP future": infected premises reported after the timepoint).

##### B.4 Posterior distributions

Posterior distributions of each inferred model parameter at each timepoint (3-, 5- and 7-weeks in) are presented by cluster: Sydney (Figure B18), CCMN (Figure B19), Tamworth (Figure B20), and Hunter Valley (Figure B21). Note that we have also included posterior distributions recovered from the full dataset (i.e., $t_{\text{today}} = t_{\text{max}} - 1$) for each cluster.

###### B.4.1 Effect of premises density on local spread

Parameters on the non-linear effect of premises density on local spread ($\xi$, $\zeta$) appeared to be the most sensitive, with posterior estimates diverging the most from their uniform priors and increased in precision through time across all clusters. Moreover, these estimates varied in both direction and strength between clusters.

Posterior estimates for $\xi$ at timepoint 3 indicated that higher density premises were much less susceptible compared to lower density premises in the Sydney cluster ($\xi = -1.46[-1.71, -1.17]$; e.g., a premises of 10 horses/ha was 29 times *less* likely to be infected than a premises with 1 horse/ha), but were more susceptible than lower density premises in the CCMN ($\xi = 0.35[-0.45, 0.68]$; e.g., a premises of 10 horses/ha was 2.2 times *more* likely to be infected than a premises with 1 horses/ha) and Hunter Valley ($\xi = 0.8[0.1, 1.1]$; e.g., a premises of 10 horses/ha was 6.3 times *more* likely to be infected than a premises with 1 horse/ha) cluster. Density had a negligible effect on premises susceptibility in the Tamworth cluster ($\xi = -0.09[-0.85, 0.58]$).

Posterior estimates for $\zeta$ at timepoint 3 indicated that higher density premises were more infectious compared to lower density premises in the Tamworth ($\zeta = 0.6[-0.82, 1.15]$) and Hunter valley ($\zeta = 0.32[-0.52, 0.65]$) cluster, where a premises of 10 times the density would be 4 and 2.1 times *more* likely to infect other premises respectively. Conversely, higher density premises were less infectious compared to lower density premises in the CCMN cluster ($\zeta = -0.58[-1.16, 0.61]$), where a premises of 10 times the density would be 3.8 times *less* likely to infect other premises. Density had a negligible effect on premises infectivity in the Sydney cluster ($\xi = -0.06[-0.44, 0.37]$).

##### B.4.2 Shape of the spatial transmission kernel

Posterior distributions of spatial kernel shape ($\psi$) diverged from their flat prior distributions in the CCMN and Tamworth clusters, but estimates were generally imprecise and did not improve through time (4.36[0.90,17.68] and 4.07[0.32,17.72] at timepoint 3 respectively). No divergence from the prior was observed in the Sydney and Hunter Valley cluster.

##### B.4.3 Background and baseline transmission rates

There was no learning (i.e., posterior distribution diverging from the prior)on background spread rate ($\alpha$) in all clusters and timepoints.

The amount of learning observed in the baseline transmission rate ($\beta_0$) appeared to positively correlate with the size of the EI cluster. Posterior estimates in the Sydney cluster were the most precise, followed by the CCMN, and in the smallest clusters (Tamworth and Hunter Valley), were very imprecise. Despite these regional differences, a trend of increasing precision in estimates and decreasing median estimates was observed across all clusters,

##### B.4.4 Effect of vaccination

Posterior distributions of the vaccine effectiveness parameter ($\theta$) did not diverge from its flat prior across all clusters and timepoints. This was also observed in subsequent model fits to the full dataset (i.e., $t_{\text{today}} = t_{\max} - 1$), though some information gain (relative to the prior) was observed only in the heavily vaccinated CCMN cluster ($\theta = 0.69 = [0.09, 0.99]$). See Section B.6 for further details.

##### B.4.5 Intra-premises transmission rate

Posterior estimates on intra-premises transmission rate ($\beta_{\text{intra}}$) diverged from its wide uniform prior ($\mathcal{U}(1, 20)$) but were generally similar across timepoints and clusters (95% CrI $\approxeq [1, 5]$). Corresponding estimates of $R0_{\text{intra}}$ are provided in Table B2, calculated from posterior estimates of $\beta_{\text{intra}}$ (i.e., $R0_{\text{intra}} = \beta_{\text{intra}} \cdot 1/\gamma_{\text{intra}}$, where $\gamma_{\text{intra}} = 1/6$).

Table B2: Estimates of $R0_{\text{intra}}$ by cluster and timepoint, calculated from posterior estimates of $\beta_{\text{intra}}$ (see text for details). Median estimates accompanied by their 95% credible intervals (in brackets) are provided.

| | 3-weeks in | 5-weeks in | 7-weeks in |
|---|---|---|---|
| Sydney | 15.98 [6.59,28.76] | 16.79 [6.54,28.74] | 15.84 [6.51,28.79] |
| CCMN | 22.78 [7.07,71.2] | 19.04 [7.24,29.3] | 16.87 [6.74,29.1] |
| Tamworth | 18.33 [6.98,29.71] | 16.53 [6.56,28.59] | 16.43 [6.56,28.77] |
| Hunter | 17.55 [6.51,29.08] | 16.51 [6.8,28.9] | 16.76 [6.63,28.87] |

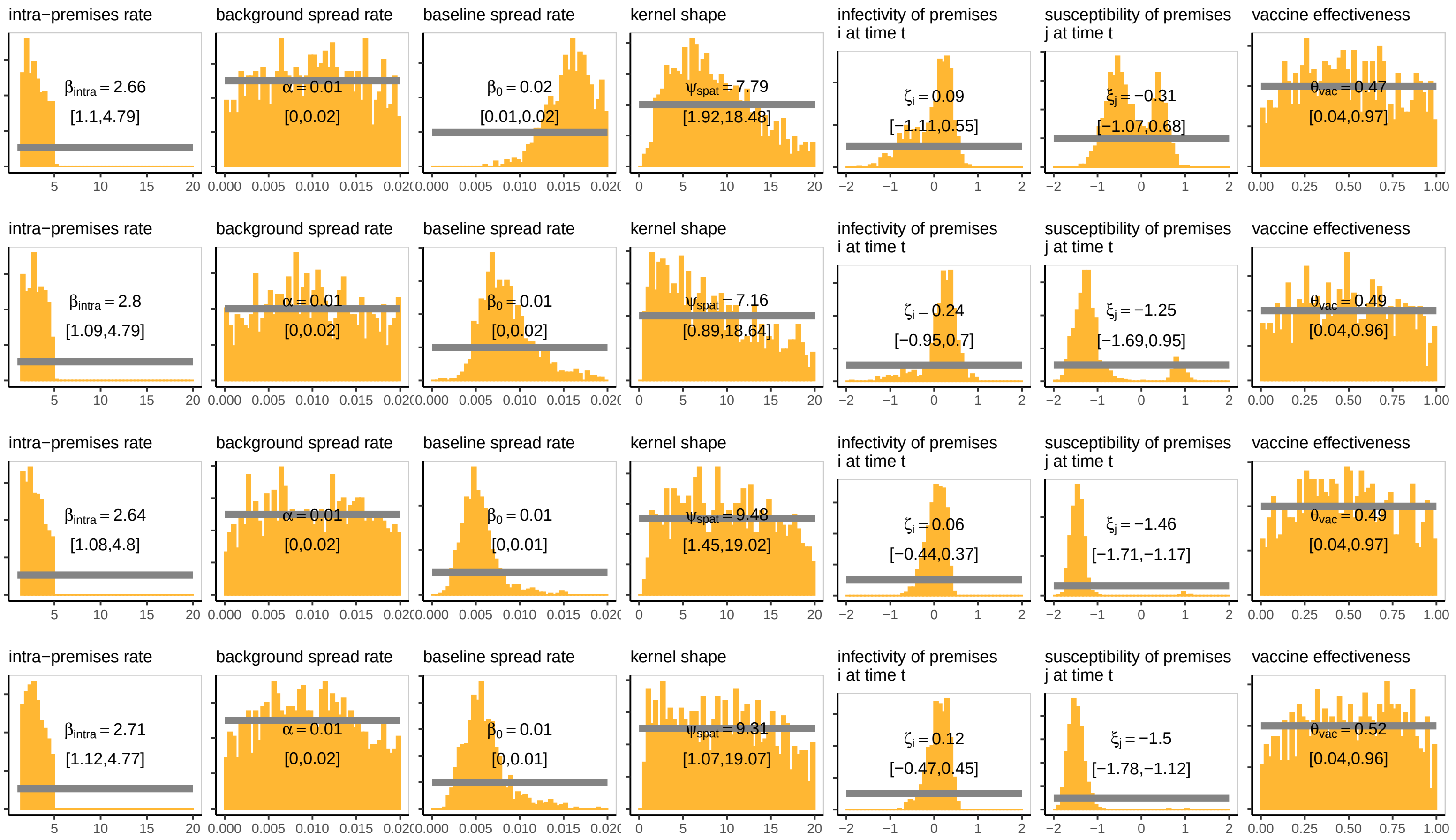

Figure B18: Prior (grey) and posterior (orange) distributions for all unknown model parameters based on data available at all three timepoints (3-, 5-, 7-weeks in) and at the end of the outbreak (bottom panel) for the *Sydney Basin region*.

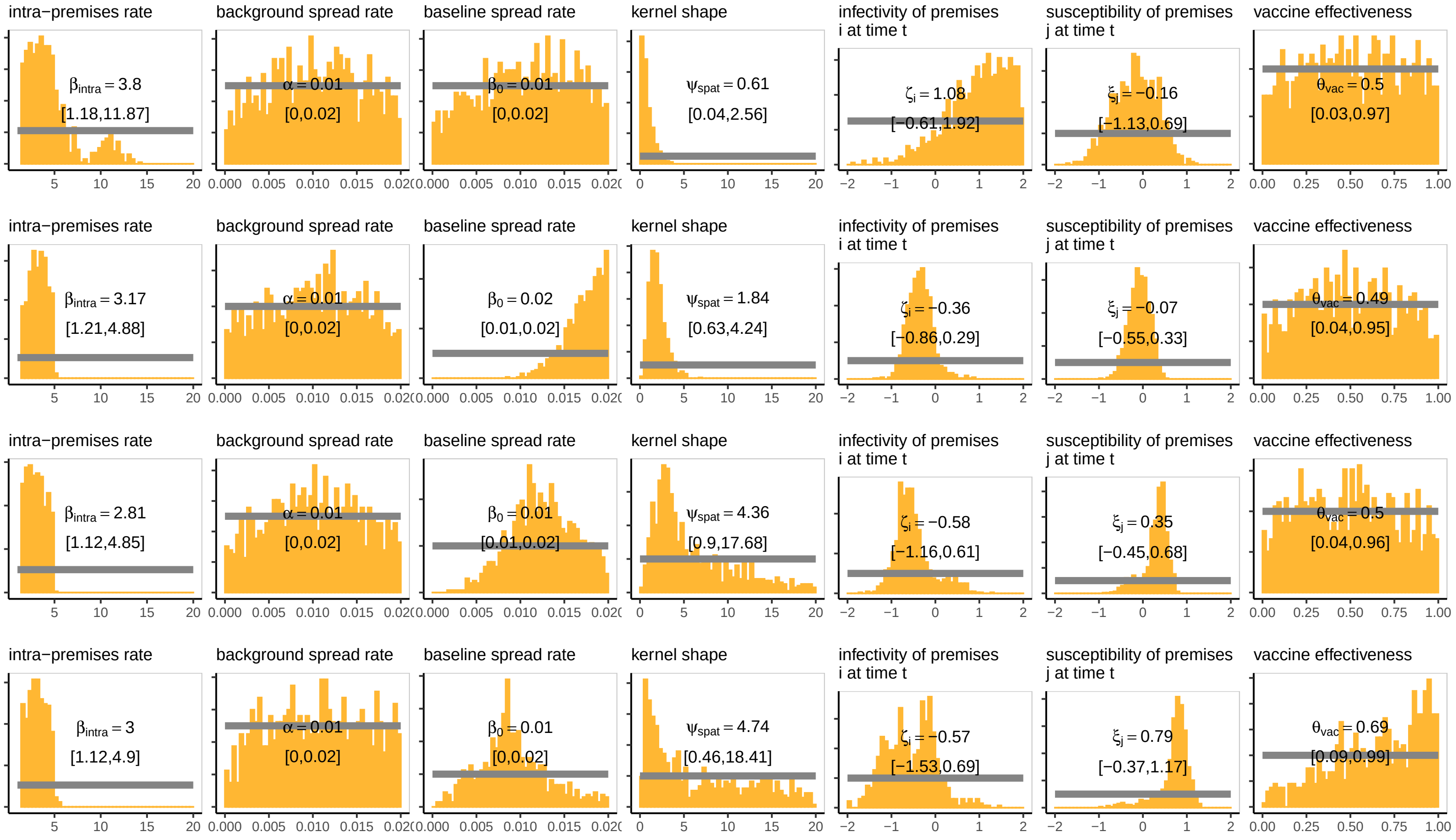

Figure B19: Prior (grey) and posterior (orange) distributions for all unknown model parameters based on data available at all three timepoints (3-, 5-, 7-weeks in) and at the end of the outbreak (bottom panel) for the *Central Coast-Maitland-Newcastle region.*

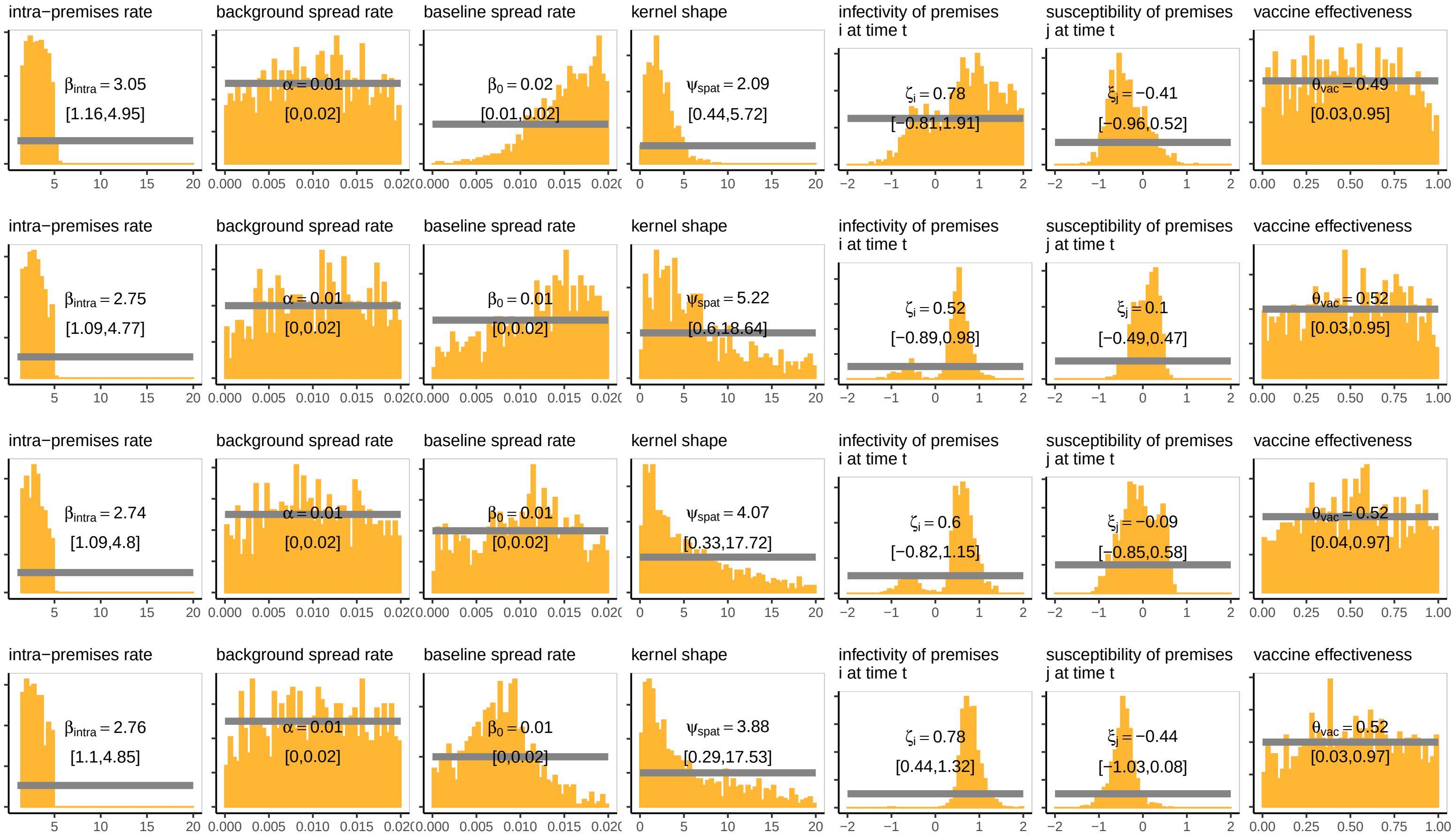

Figure B20: Prior (grey) and posterior (orange) distributions for all unknown model parameters based on data available at all three timepoints (3-, 5-, 7-weeks in) and at the end of the outbreak (bottom panel) for the *Tamworth region*.

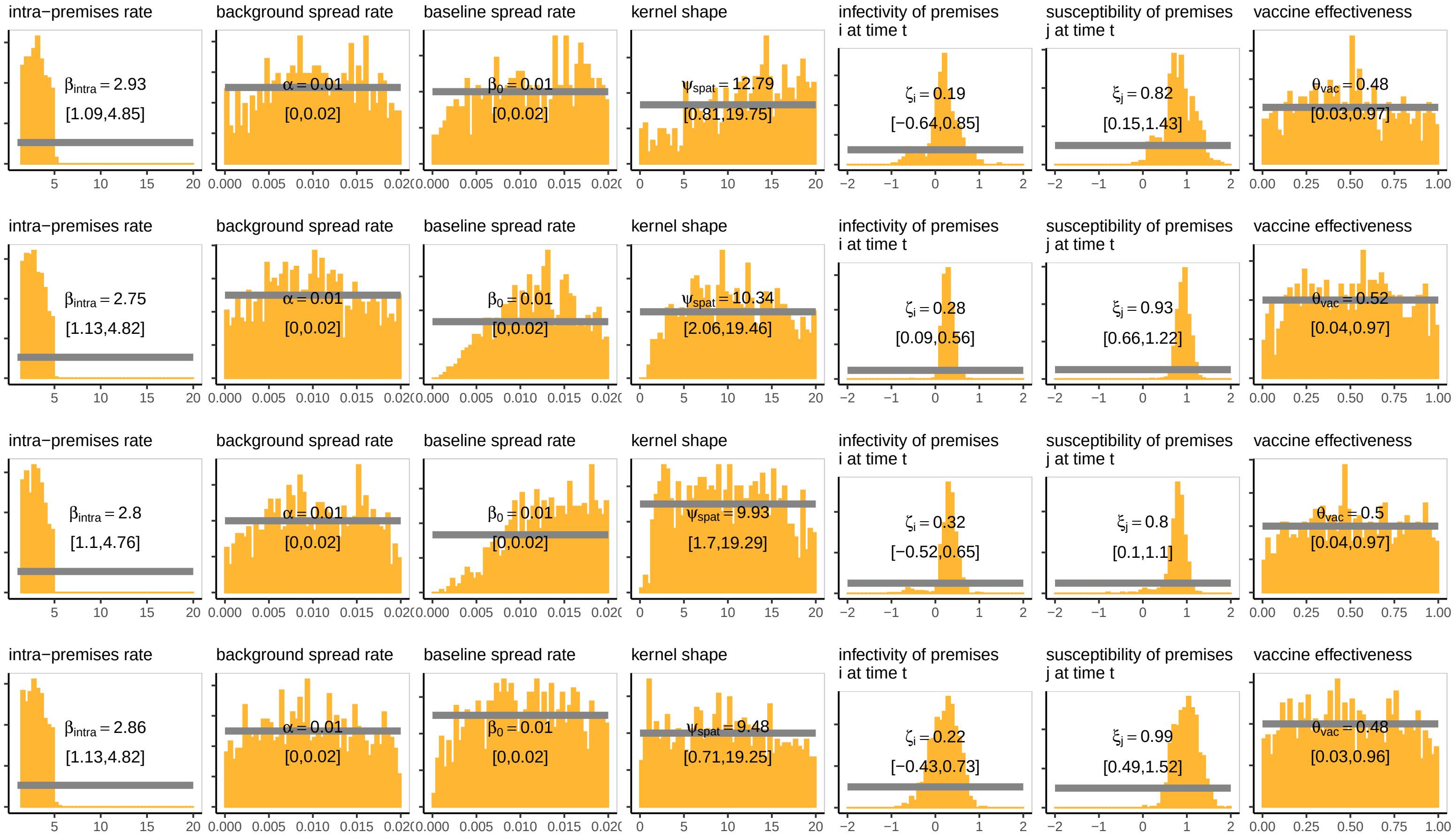

Figure B21: Prior (grey) and posterior (orange) distributions for all unknown model parameters based on data available at all three timepoints (3-, 5-, 7-weeks in) and at the end of the outbreak (bottom panel) for the *Hunter Valley region*.

#### B.5 Forecast skill

Figure B22 shows forecast skill by lead time for each of the four EI outbreak clusters.

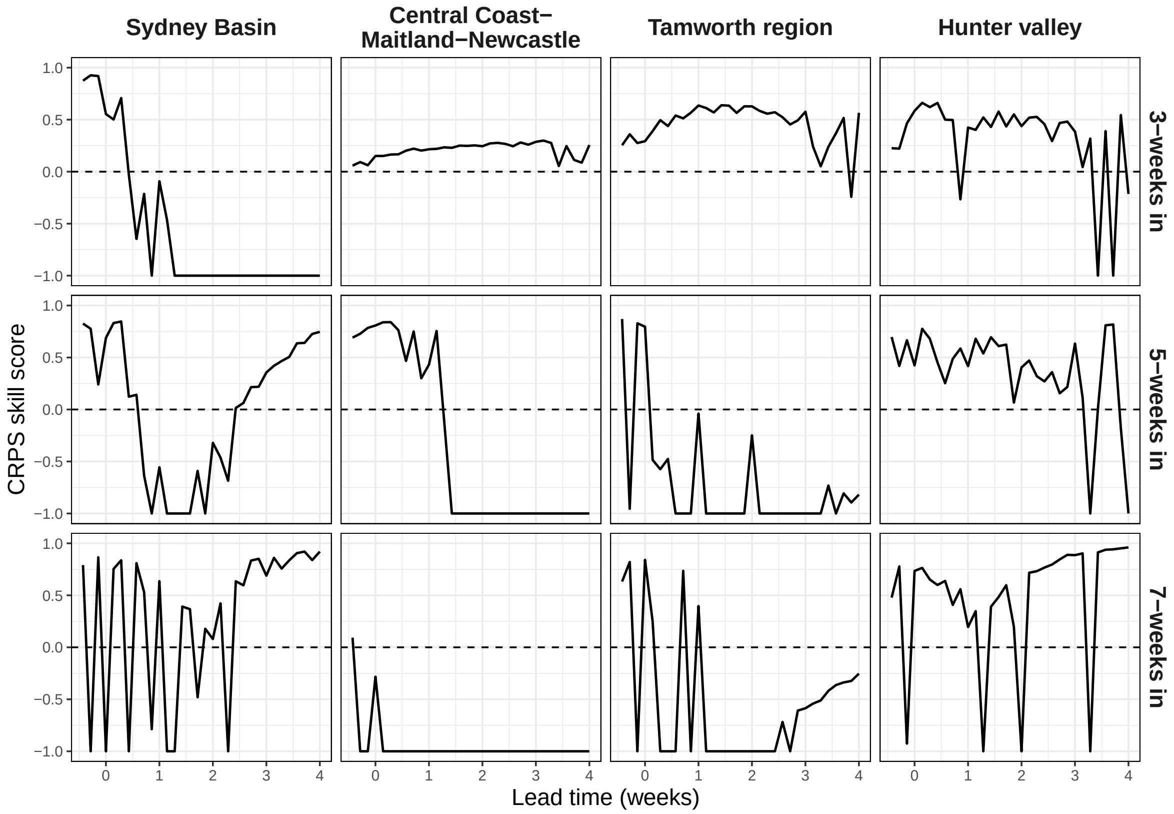

Figure B22: Forecast skill of EI daily case counts by lead time for four outbreak clusters in Australia, shown separately for each time point (right-hand axis labels). Positive skill scores indicate that the forecasts outperformed the naive benchmark, with a maximum possible score of one (i.e., perfect accuracy and no uncertainty). Negative skill scores indicate underperformance relative to the naive benchmark, with a mininum score of $-\infty$. Skill scores below -1 were truncated.

We also considered forecast skill as a function of the number of cases ultimately recorded on each day in each cluster (comprising 384 observations). We divided the daily case counts into four bins: 0 cases (n = 36, 9.4%); 1–4 cases (n = 78, 20.3%); 5–9 cases (n = 69, 18.0%); 10–29 cases (n = 129, 33.6%); and $\geq$ 30 cases (n = 72, 18.8%). There appeared to be no consistent trend between forecast skill and daily case numbers across the four clusters (Figure B23).

Spatial forecast skill generally improved after the first timepoint (3-weeks in), and tended to

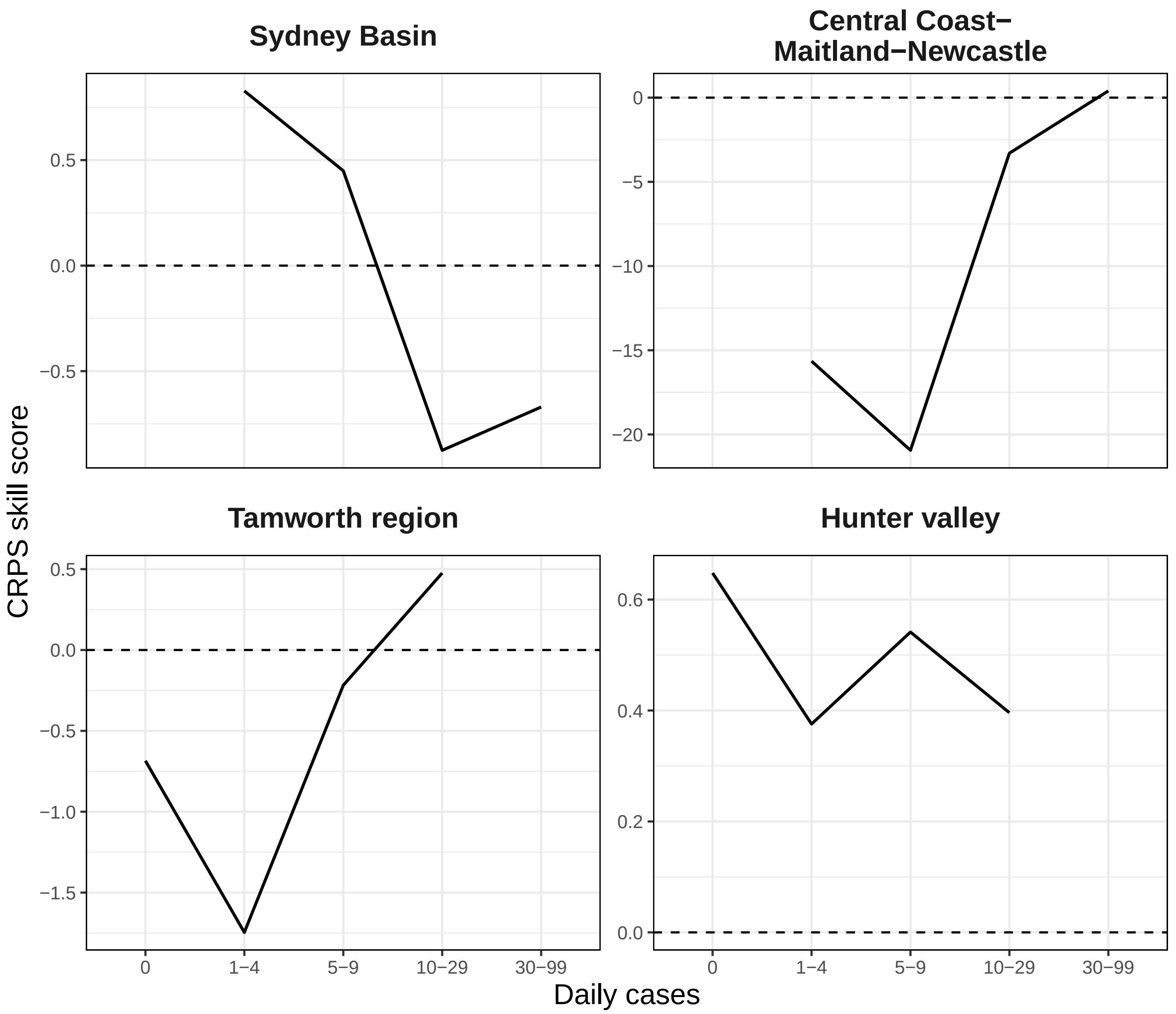

Figure B23: CRPS skill scores binned by the number of cases ultimately recorded for the day of prediction shown separately for each outbreak cluster.

be higher for 2-weeks ahead projections (Figure B24).

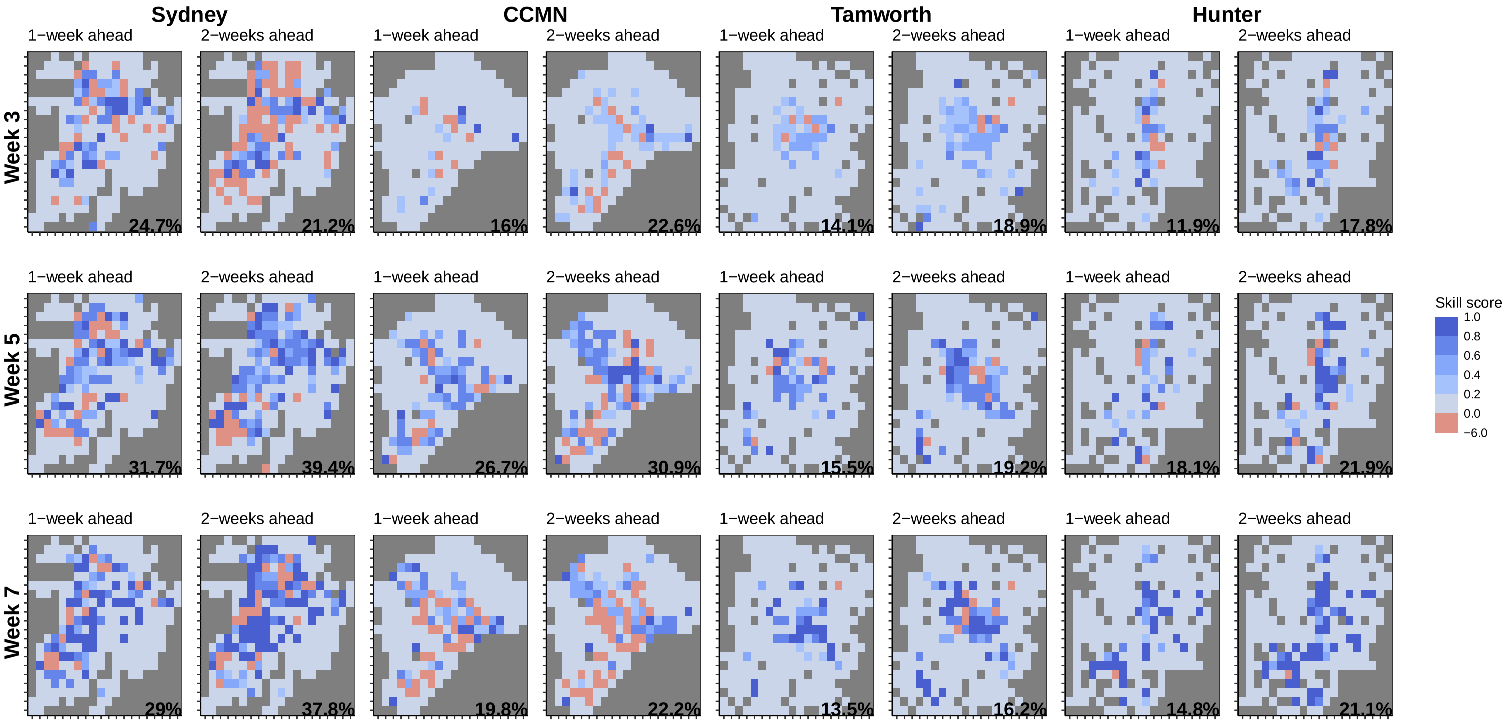

Figure B24: Spatial forecast skill for 1- and 2-week ahead case counts for each cluster and timepoint.

#### B.6 Vaccination

We provided an initial demonstration through our 2007 EI outbreak case study where we evaluated the impact of the implemented vaccination policy through our vaccine effectiveness parameter $\theta$ (Section B.4.4).

To have a noticeable effect at the population-level, the vaccination rate would have to outpace the infection rate (Probert 2018); though in local clusters within disease-free premises, they can have a protective effect. The lack of information gained relative to the prior for $\theta$ suggests that the implemented vaccination policy did little to contain outbreak, corroborating the findings of other analyses of the 2007 EI outbreak (Garner et al. 2011; Firestone 2012). It did not matter that substantial vaccinations were occurring because they occurred in the peripheral regions of each cluster and when the outbreak was already dying out (Figure B25, Figure B26, Figure B27, Figure B28). Furthermore, peak immunity took around 14 days to reach (Edlund Toulemonde et al. 2005). Though, the model fit to the full dataset provided some indication that vaccinations had some protective effect only in the CCMN cluster and could have prevented a second wave from occuring, possibly due to a large proportion of the cluster being vaccinated. These findings are preliminary and should be investigated further in future work.

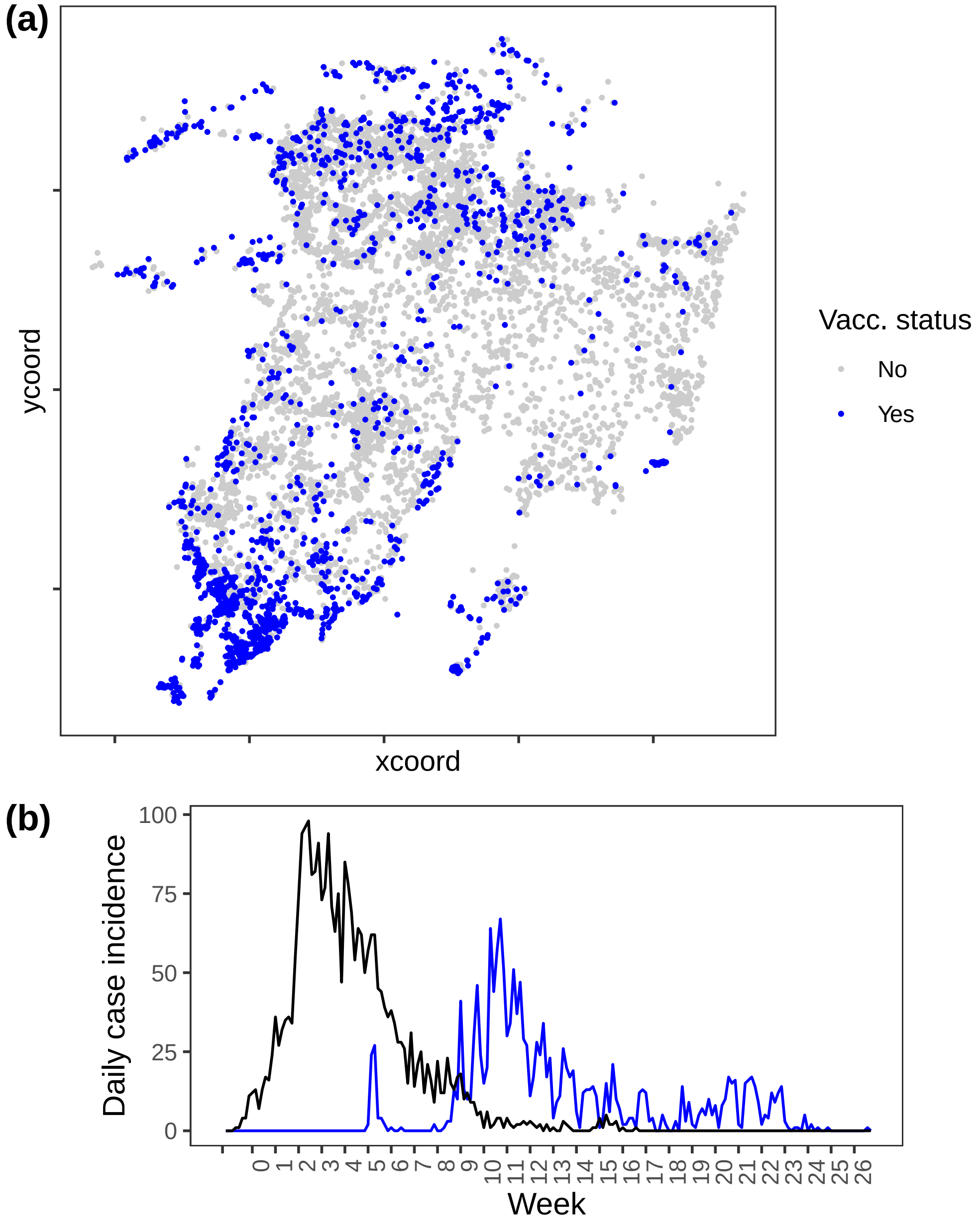

Figure B25: Vaccinations in the *Sydney region*.

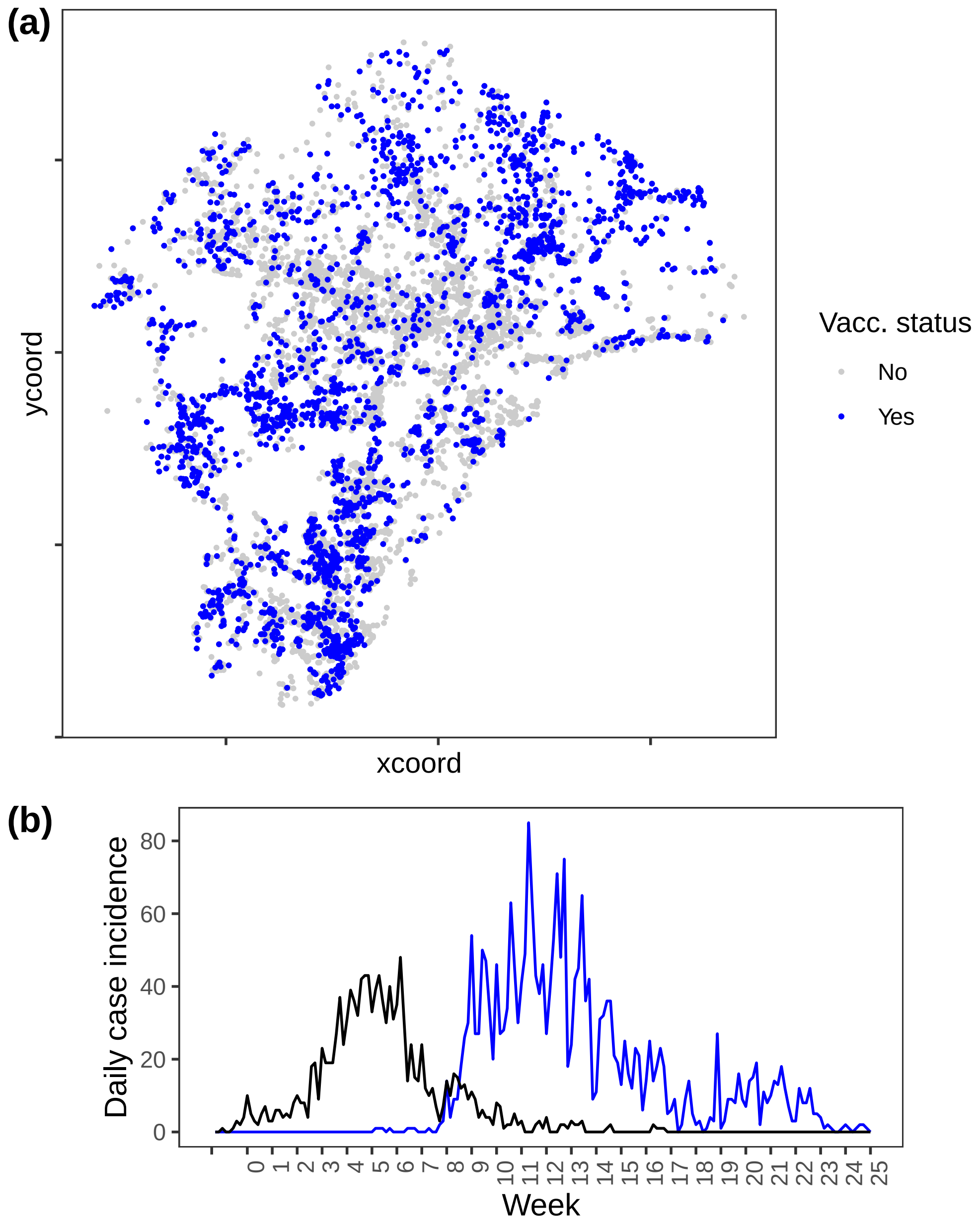

Figure B26: Vaccinations in the *Central Coast-Maitland-Newcastle region*.

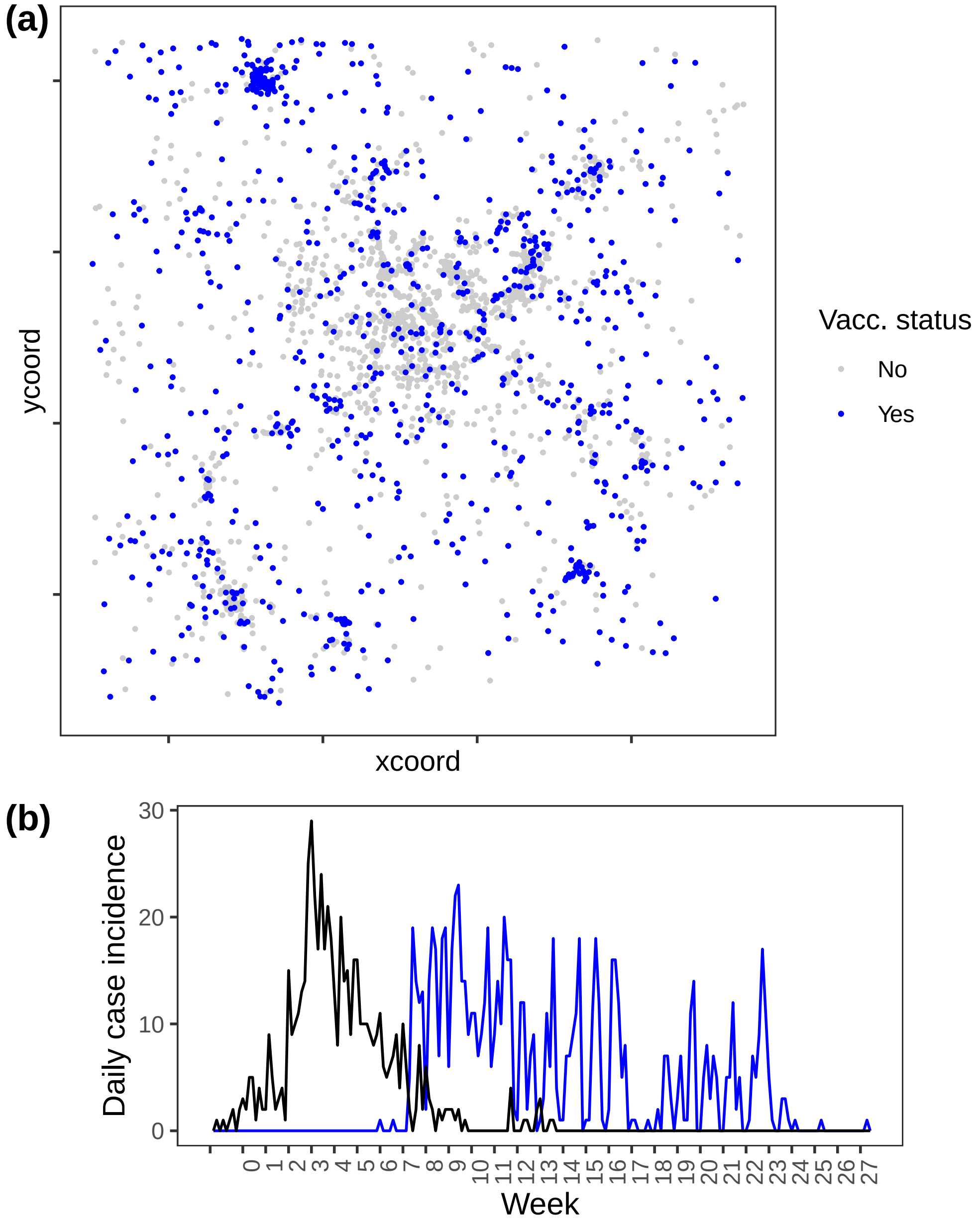

Figure B27: Vaccinations in the *Tamworth region*.

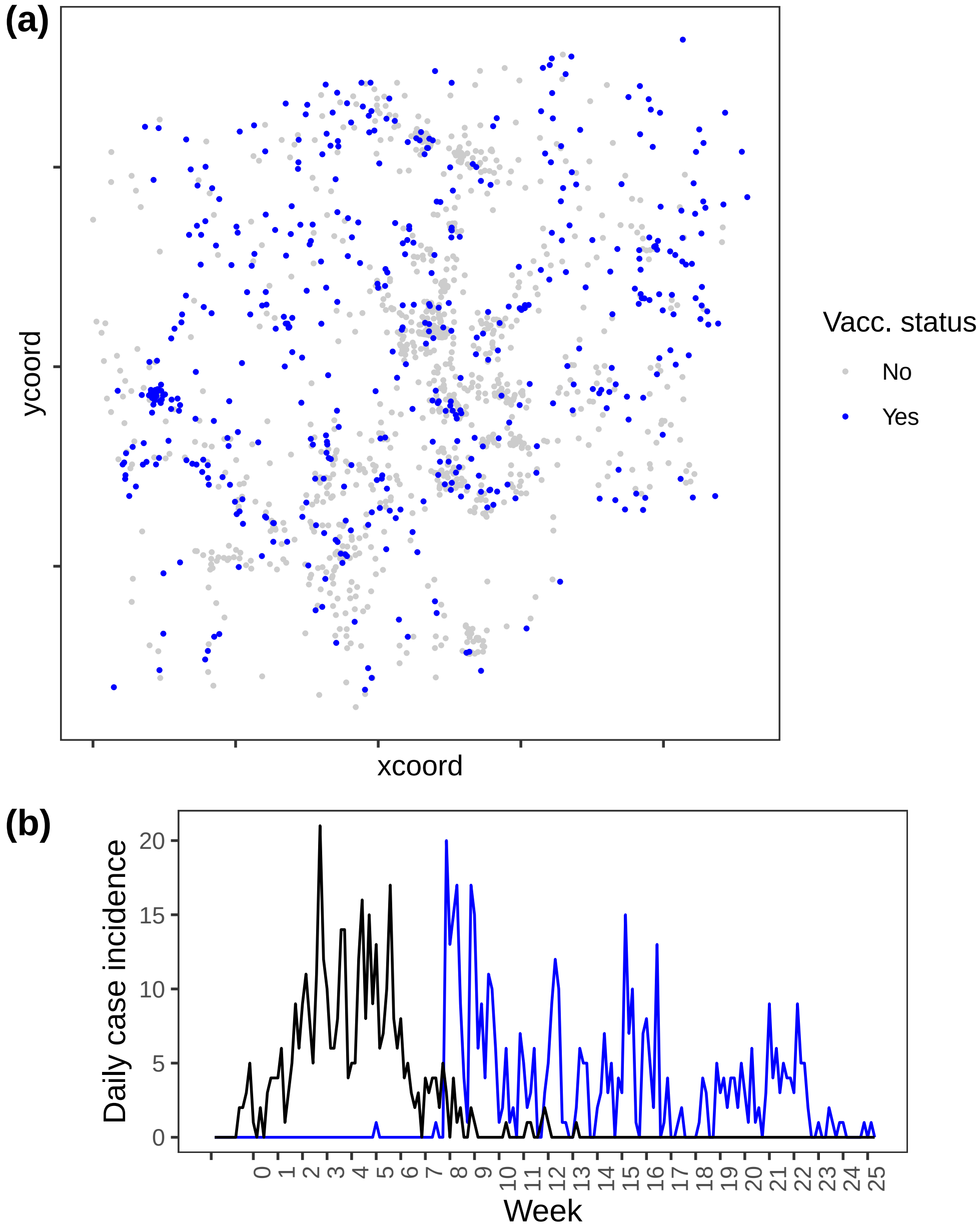

Figure B28: Vaccinations in the *Hunter Vallery region.*
